# Into the Wild: A novel wild-derived inbred strain resource expands the genomic and phenotypic diversity of laboratory mouse models

**DOI:** 10.1101/2023.09.21.558738

**Authors:** Beth L. Dumont, Daniel Gatti, Mallory A. Ballinger, Dana Lin, Megan Phifer-Rixey, Michael J. Sheehan, Taichi A. Suzuki, Lydia K. Wooldridge, Hilda Opoku Frempong, Gary Churchill, Cathleen Lutz, Nadia Rosenthal, Jacqueline K. White, Michael W. Nachman

## Abstract

The laboratory mouse has served as the premier animal model system for both basic and preclinical investigations for a century. However, laboratory mice capture a narrow subset of the genetic variation found in wild mouse populations. This consideration inherently restricts the scope of potential discovery in laboratory models and narrows the pool of potentially identified phenotype-associated variants and pathways. Wild mouse populations are reservoirs of predicted functional and disease-associated alleles, but the sparsity of commercially available, well-characterized wild mouse strains limits their broader adoption in biomedical research. To overcome this barrier, we have recently imported, sequenced, and phenotyped a set of 11 wild-derived inbred strains developed from wild-caught *Mus musculus domesticus*. Each of these “Nachman strains” immortalizes a unique wild haplotype sampled from five environmentally diverse locations across North and South America: Saratoga Springs, New York, USA; Gainesville, Florida, USA; Manaus, Brazil; Tucson, Arizona, USA; and Edmonton, Alberta, Canada. Whole genome sequence analysis reveals that each strain carries between 4.73-6.54 million single nucleotide differences relative to the mouse reference assembly, with 42.5% of variants in the Nachman strain genomes absent from classical inbred mouse strains. We phenotyped the Nachman strains on a customized pipeline to assess the scope of disease-relevant neurobehavioral, biochemical, physiological, metabolic, and morphological trait variation. The Nachman strains exhibit significant inter-strain variation in >90% of 1119 surveyed traits and expand the range of phenotypic diversity captured in classical inbred strain panels alone. Taken together, our work introduces a novel wild-derived inbred mouse strain resource that will enable new discoveries in basic and preclinical research. These strains are currently available through The Jackson Laboratory Repository under laboratory code *NachJ*.

## INTRODUCTION

Inbred mouse strains have served as the workhorses of mammalian genetics for nearly a century (1). Standardized inbred strain backgrounds ensure experimental reproducibility across labs and experiments, provide the backbone for mechanistic investigations into gene and pathway function, and enable testing of genetically identical cohorts across different treatments, exposures, and perturbations. Inbred strains also provide platforms for community resource development, including comprehensive gene knockout panels (2,3) and phenome resources (4).

The classical inbred (CI) mouse strains were developed from a small number of founder mice purpose-bred by mouse fanciers for traits of interest in the early 1900s (5,6). As a result, the CI strains capture a limited subset of the genetic variation found in wild mouse populations (7). Many CI strains have inherited large stretches of their genome identical by descent, such that pairwise strain comparisons yield numerous genomic regions where segregating variation is reduced to or near zero (8,9). Furthermore, due to their unique origins and history of selective breeding, the complex architecture of trait variation in inbred strains may not faithfully model the genetic organization of phenotypic variation in human populations (10).

Wild mouse populations harbor numerous predicted functional and disease-associated alleles, the majority of which are not present in classical inbred mouse strains and have therefore never been experimentally tested in the laboratory (8,11). Thus, wild mice present an untapped opportunity for advancing new biomedical research discoveries (7,8,11). In contrast to the history of intense artificial selection and substantial introgression between subspecies that has molded the genomes of classical laboratory strains, the genetic diversity observed in wild mice reflects the interplay of selection, genetic drift, mutation, and migration, mirroring the natural population genetic processes that have sculpted the contemporary landscape of human genomic diversity. Wild mice may therefore better approximate the diversity and organization of functional genetic variation in human populations than CI strains, including genetic variation influencing responses to dietary challenges, pharmaceutical interventions, and toxin exposures.

Despite these potential advantages, multiple challenges stand in the way of using of wild-caught mice directly in biomedical research. Trapping wild house mice is laborious, especially if one is interested in assembling a large sample of unrelated individuals. Further, wild mice are genetically unique, preventing experimental designs that require controlled genetic backgrounds. In addition, wild mice exhibit phenotypic variation as a result of differences in age, reproductive history, health status and other, typically unknown, environmental exposures. Finally, wild mice are also vectors for numerous pathogens that pose a threat to human health and the health status of laboratory mouse colonies.

Wild-derived inbred mouse strains (WDIS) present a powerful intermediary between CI and wild mice. WDIS are developed from wild-caught mice that are brother-sister mated in a laboratory environment for >20 generations, thereby immortalizing a single haplotype from the wild in an inbred state. Thus, WDIS combine the reproducibility and fixed genetic background of inbred mouse models with the increased diversity present in wild mouse populations. WDIS have been featured as founders to current mouse diversity populations such as the Collaborative Cross (CC) and Diversity Outbred (DO; (12,13)), strategically utilized in gene mapping studies to introduce increased diversity (14–18), and profiled in immunological (19), metabolic (20), and reproductive studies (21–23). Indeed, the majority of quantitative trait loci (QTL) identified in mapping studies in the CC and DO are attributable to the allelic effects of one or more of the three wild-derived founder haplotypes (24–27).

Despite their realized power, only a modest number of wild-derived inbred strains are commercially available (**Supplementary Table 1**). Of these, many are poor breeders, are maintained at low (or undocumented) health status, or have few associated genomic resources. These considerations present notable obstacles to their widespread use in biomedical research. Furthermore, whereas the genomes of CI strains are largely derived from one house mouse subspecies (*M. m. domesticus*) (6), many WDIS are representatives of alternative house mouse subspecies that exhibit variable degrees of reproductive isolation from *M. m. domesticus*. Crosses between these WDIS and CI strains often expose multilocus incompatibilities linked to hybrid sterility (28,29), an outcome that further limits their practical utility in genetic studies.

Recently, we developed a panel of ∼25 wild-derived inbred strains from wild-caught *M. m. domesticus* from five locations across North and South America: Saratoga Springs, New York, USA; Gainesville, Florida, USA; Manaus, Brazil; Tucson, Arizona, USA; and Edmonton, Alberta, Canada (**Figure 1**). These sampling locations are defined by distinct ecosystems and climates, including tropical rainforest (Manaus), desert (Tucson), temperate forest (Saratoga Springs), prairie (Edmonton), and wetland (Gainesville) habitats. As a result, wild mice from these regions have been subject to distinct selective pressures and have evolved unique morphological and physiological adaptations to their environments (30–33). Recent work has also discovered dramatic transcriptomic differences between populations, revealing divergence at the functional genomic level (32). Taken together, data from wild mice sampled from these five geographic regions portend the rich potential for their descendant inbred strains to serve as powerful new mouse models of resistance and susceptibility to numerous diseases, phenotypes, and conditions relevant to human health.

**Figure 1.**
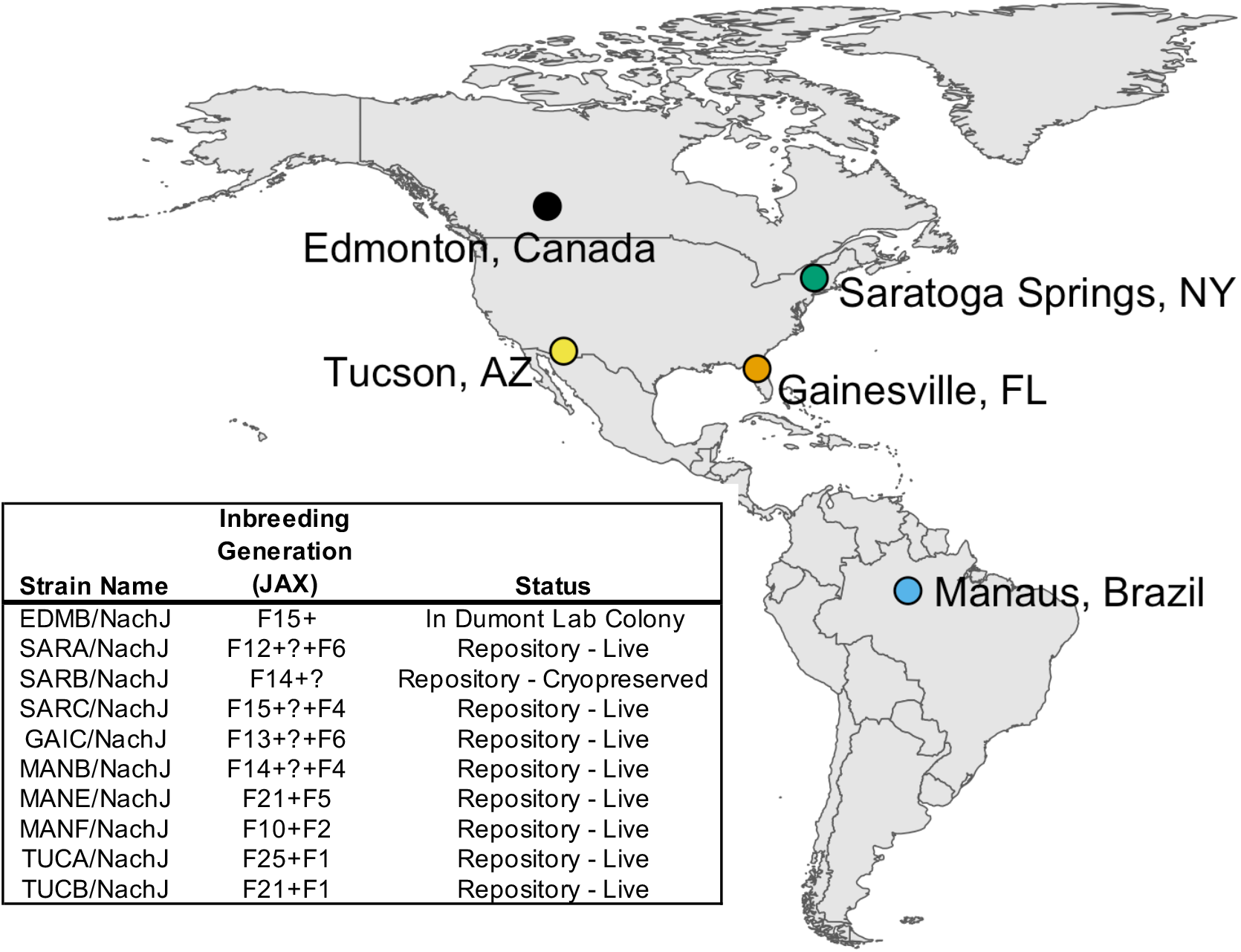
Map of sample locations and summary of inbred strains imported to the JAX Repository. Inbred strains were developed from wild-caught mice trapped at 5 geographic sample locations across North and South America: Manaus, Brazil (MANB/NachJ, MANE/NachJ, MANF/NacJ); Saratoga Springs, New York, USA (SarA/NachJ, SARB/NachJ, SARC/NachJ); Gainesville, Florida, USA (GAIC/NachJ); Tucson, Arizona, USA (TUCA/NachJ, TUCB/NachJ); and Edmonton, Alberta, Canada (EDMB/NachJ). Strain GAIA/NachJ was successfully imported but has been since discontinued from the JAX Repository due to poor breeding and is not listed. Inbreeding generation numbers are imprecise owing to the use of inter-generational crosses in JAX’s Importation facility to expedite colony expansion and oocyte harvests. This ambiguity is denoted by a “?” in the inbreeding generation supplied in the strain table.

From 2019-2022, we imported a representative subset of 11 wild-derived inbred strains from the broader parent Nachman panel to The Jackson Laboratory (JAX), including at least one inbred line per geographic location (**Supplementary Table 2**). Here, we introduce this novel diverse mouse strain resource, including the extent of genomic and phenotypic diversity across these strains and relationship to CI strains. Our strain survey emphasizes the collective potential of these Nachman strains to advance biomedical discoveries into multiple trait domains, systems genetics analysis, and fundamental principles of evolutionary biology.

## RESULTS

### Generating the Nachman Panel

Wild house mice were caught in 2013 from five geographic locations (**Figure 1**). Animals were transported to UC Berkeley, and mice from each geographic region were randomly paired to create inbred lines which were propagated through brother-sister mating for at least 10 generations. Initially, ∼10 independent lines were established from each location. No attempts were made to rescue lines that exhibited infertility due to inbreeding depression, and half of the initiated lines eventually became extinct. At the time of writing, 25 of the initiated lines remain in Dr. Nachman’s strain holdings at UC Berkeley.

A subset of 20 strains were selected for importation to the JAX Repository (**Figure 1**). Of these strains, 11 were successfully rederived via *in vitro* fertilization and embryo transfer to a pseudopregnant dam and integrated into production breeding colonies. Reasons for rederivation failure were complex and variable, ranging from poor breeding, failure to recover sufficient numbers of oocytes for IVF, no live births following multiple embryo transfers, and accidental strain contamination (**Supplementary Table 2**). One strain (GAIA/NachJ) was subsequently terminated due to poor breeding performance.

### Breeding Performance

Many wild-derived inbred strains breed poorly, a consideration that limits their practical utility in biomedical research. Our strategic propagation of only the best performing inbreeding lineages derived from each wild-caught founder pair should ensure that resulting inbred strains are robust breeders. Indeed, the inbred Nachman lines breed reliably across two independent animal facilities (**Figure 2**). Fewer than 25% of established matings are non-productive and average weaned litter sizes for most strains are between 4-6 pups.

**Figure 2.**
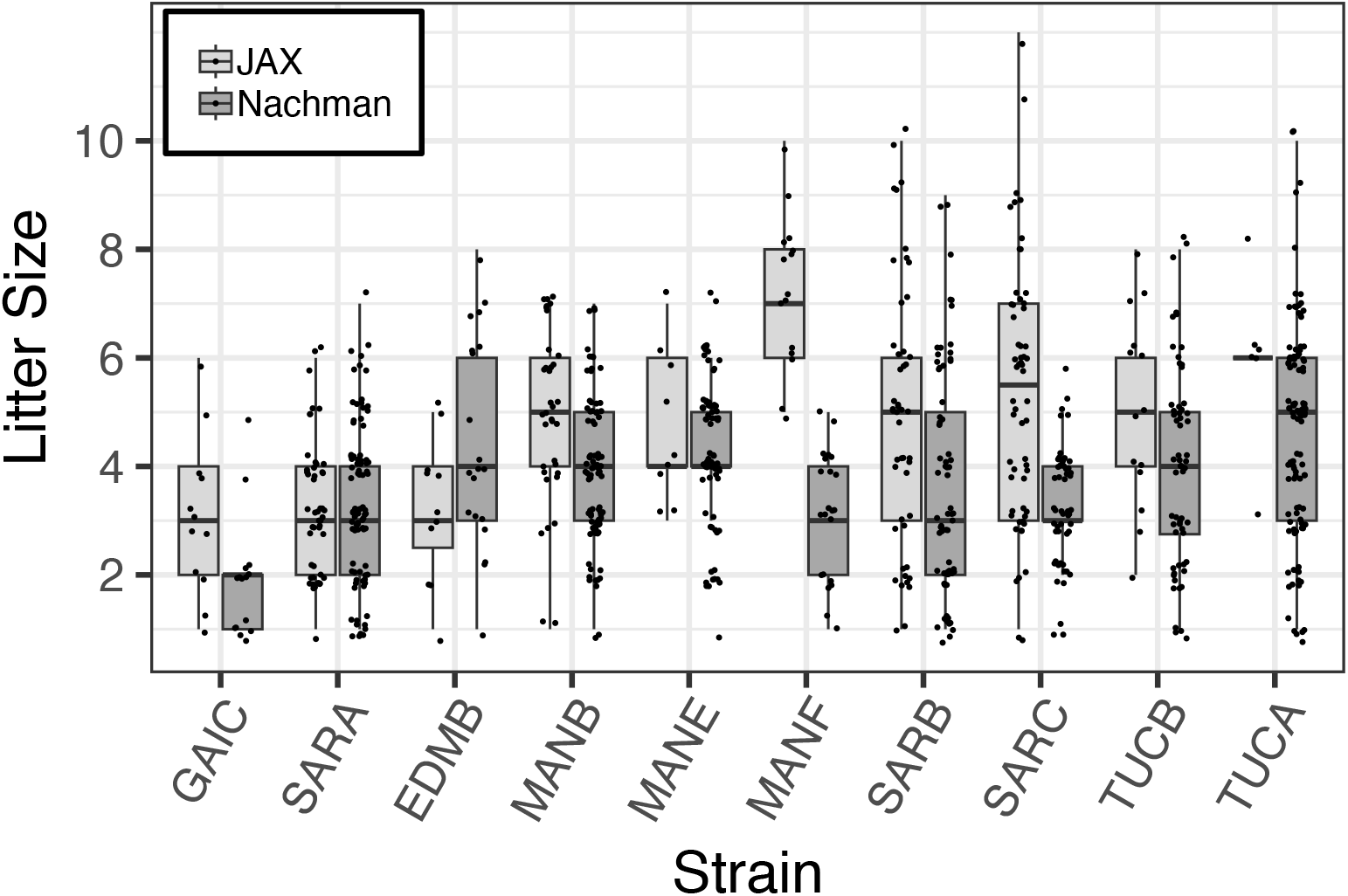
Distribution of weaned litter sizes across 10 strains in the Nachman wild-derived inbred strain panel. Sizes of litters born at JAX and in the Nachman Lab at UC Berkeley are presented as standard box plots, with box width defining the inter-quartile range and the thick line bisecting each box denoting the median. Litter sizes are jittered for ease of visualization.

While comprehensive diallele crosses have not yet been performed, there are no signs of F1 sterility or reduced fertility in the few F1 hybrids between independent Nachman strains that have been generated to date (**Supplementary Table 3**). In contrast, testis weight and sperm density in F1 hybrids exceed fertility metrics quantified in the inbred parental lines and classical inbred strains, revealing hybrid vigor (**Supplementary Figure 1; Supplementary Table 3**). Efforts to comprehensively profile reproductive traits in the inbred Nachman strains and their derivative F1 hybrids are on-going, and we expect our future results to strengthen empirical support for high F1 fertility. These preliminary findings fall in contrast to observations from diallele crosses involving inbred founder strains from the DO and CC (34), which feature strains derived from reproductively isolated subspecies. Many incipient inbred CC strains were lost due to male infertility (35) and many surviving strains exhibit low breeding performance, male infertility, and extreme sex ratio distortion (36). Thus, we anticipate that strains in the Nachman wild-derived strain panel comprise a shelf-stable mouse diversity resource well-suited for genetic crosses.

### Cytogenetic Characterization of Nachman Strain Karyotypes

Wild *M. m. domesticus* populations harbor frequent Robertsonian chromosomal translocations that give rise to considerable karyotypic diversity across Europe (37). Hybrid mice from crosses between different karyotypic races exhibit reduced fitness owing to altered chromosome dynamics and inefficient chromosome segregation during meiosis (37–40). To confirm the absence of large-scale karyotypic alterations or changes in chromosome number among strains, we generated meiotic cell spreads from each inbred Nachman line and evaluated karytotype and meiotic chromosome pairing by fluorescent microscopy. **Supplementary Figure 2**. These cytogenetic analyses are on-going, but none of the five Nachman strains evaluated to date carry Robertsonian translocations (EDMB, SARA, SARB, MANB, and TUCB). Each of these five strains exhibits the standard 2n=40 all-acrocentric karyotype.

### Genomic Diversity in the Nachman Panel

To assess the extent of genomic diversity in this new strain resource, we sequenced the whole genomes of a representative male from each strain to moderate coverage using PacBio HiFi sequencing technology (∼10x coverage per strain; **Table 1**). Average read lengths exceeded 10kb for all strains (range: 10.217 – 14.957kb), with >96.67% of sequenced bases exceeding a quality score of 30 (**Supplementary Table 4**). Sequenced reads were mapped to the GRCm39 reference genome assembly (41) and subject to single nucleotide variant (SNV) calling using DeepVariant (42,43). Each strain harbors between 4.73-6.54 million fixed SNV differences relative to the C57BL/6J-based reference, with >2.75M SNVs distinguishing any pair of strains (**Figure 3A**; corresponding to a minimum of ∼1 SNP every ∼1 kb). Although sequenced mice have undergone a modest number of inbreeding generations, within strain heterozygosity is low (*π* < 0.0006 versus ∼0.0017 in wild-caught *M. m. domesticus*) and aligns with theoretical expectations for the number of inbreeding generations (**Supplementary Figure 3**).

**Figure 3.**
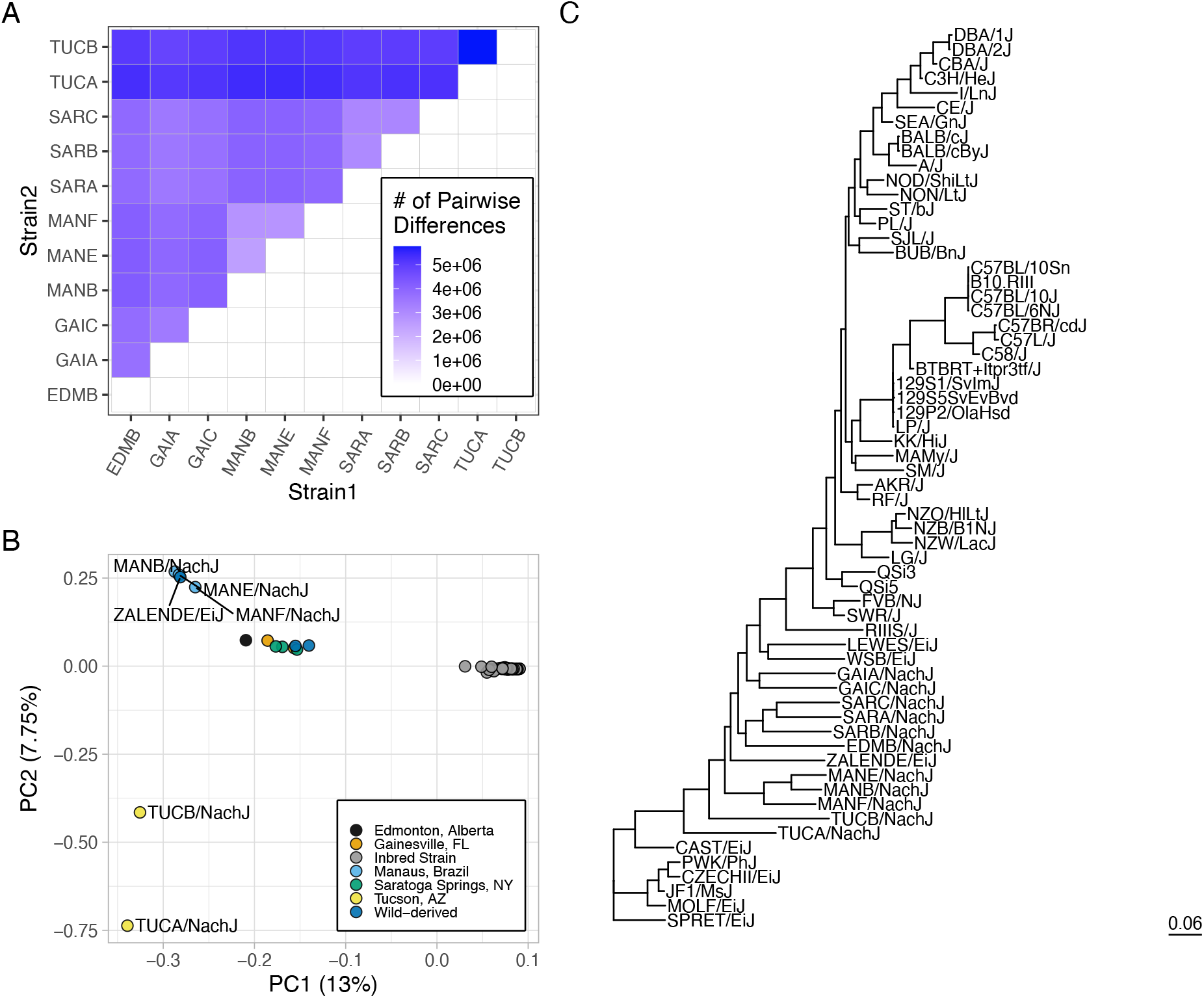
A high level of genetic diversity in the Nachman strains. (A) Heatmap displaying the number of pairwise SNP differences between Nachman lines. Counts derive from the autosomal genome fraction only and exclude unplaced contigs. (B) PCA analysis of autosomal genetic diversity partitions the Nachman lines by sample location and isolates Nachman strains from the classical inbred mouse strains. (C) Maximum likelihood phylogenetic tree constructed from SNPs on chr19. The classical inbred strains form a single clade nested within the diversity sampled by the Nachman strains.

We performed joint SNV calling with the Nachman lines and 51 classical inbred mouse strains previously sequenced by the Mouse Genomes Project (46). Of the 16,071,877 autosomal SNVs observed in the Nachman strains, 6,836,536 variants (42.5%) are absent from classical inbred mouse strains (46). We constructed a maximum likelihood phylogenetic tree from SNPs on chr19 to assess relationships between existing mouse strains and strains in this new wild-derived inbred strain resource. The classical laboratory strains present as a single clade nested within the diversity sampled in the Nachman panel (**Figure 3C**). Branch lengths are notably longer for the wild-derived inbred strains than for the classical inbred strains, reflecting the greater diversity in the former. These findings are confirmed by a principal component (PC) analysis, which reveals a single cluster of points corresponding to the classical inbred strains (**Figure 3B**), with the Nachman strains and other wild-derived inbred strains well-separated in PC coordinate space. Nachman strains derived from a common locale are more genetically similar than strains from divergent locations, although strains from Gainesville, Edmonton, and Saratoga Springs are minimally separated along PC1-PC5 (41.53% of the variance). PC dimensions 6-8 provide separation across these geographic sample locations (**Supplementary Figure 4**).

We next annotated variants according to likely functional impact using the Ensembl variant effect predictor. Variant counts for different functional categories are provided in **Supplementary Table 5**. Overall, the Nachman strain panel harbors 1,976 SNPs predicted to be highly deleterious, with 1,319 of these variants not observed in classical inbred laboratory strains. These highly deleterious variants are enriched in genes with biological roles in sensory perception and G protein-coupled receptor signaling (**Supplementary Table 5**). The Nachman strains also harbor predicted loss-of-function alleles at genes implicated in human disease. For example, the TUCA/NachJ and TUCB/NachJ lines harbor a premature stop variant in *Eif2b1* that is a predicted target of nonsense-mediated decay (NMD; chr5:124716942). *Eif2b1* encodes a subunit of the eukaryotic translation initiation factor eIF2B, which is essential for protein synthesis. Mutations in this gene have been linked to leukoencephalopathy with vanishing white matter and ovarian failure in humans (47,48). Similarly, a predicted NMD variant in *Yeats2* (chr16:20028820, stop-gain mutation) is present in GAIC/NachJ, TUCA/NachJ, and TUCB/NachJ. Mutations in this gene cause myoclonic epilepsy in humans (49). Strains MANB/NachJ, MANF/NachJ, TUCA/NachJ, and TUCB/NachJ carry a predicted loss-of-function splice-donor variant in *Ifnar1* (chr16:91302893). Humans with mutations in this gene have immunodeficiency-106 and exhibit hyperinflammatory responses to some vaccines (50).

Taken together, our analyses indicate that the Nachman strains harbor considerable genetic diversity that is not captured in existing inbred mouse strain panels. A subset of this variation is likely functional, establishing the prediction of broad phenotypic diversity among these strains and their phenotypic divergence from classical inbred mouse strains.

### Structural diversity in the Nachman Panel

Structural variants (SVs) are important contributors to phenotypic diversity, including disease risk and incidence. Prior work has suggested that house mouse genomes are burdened by higher rates of structural mutation than human genomes (51,52), leading us to posit the presence of abundant, potentially functional SVs within the Nachman strain genomes.

Using an ensemble approach to minimize false positive SV calls (see Methods), we identified 53,639 autosomal structural variants across the 11 sequenced Nachman strain genomes (13,481 deletions, 38,246 insertions, 1829 duplications, and 83 inversions). Of the 13,481 deletions discovered in the Nachman strains, 5,817 were not previously detected by the Sanger Genomes Project (43.1%; requiring 75% reciprocal overlap). Nachman strain genomes each contain 8.26 Mb - 11.14 Mb of sequence that is absent in the mm39 C57BL/6J-based reference genome (**Figure 4A**), with 9.90 Mb - 12.16 Mb of sequence in the reference genome absent from any given Nachman strain (**Figure 4B**). The modest genome coverage of the Nachman lines may lead to an appreciable number of missed SVs (i.e., false negatives) in these genomes, implying that SVs potentially have a greater collective impact on these genomes than suggested by the numbers presented here.

**Figure 4.**
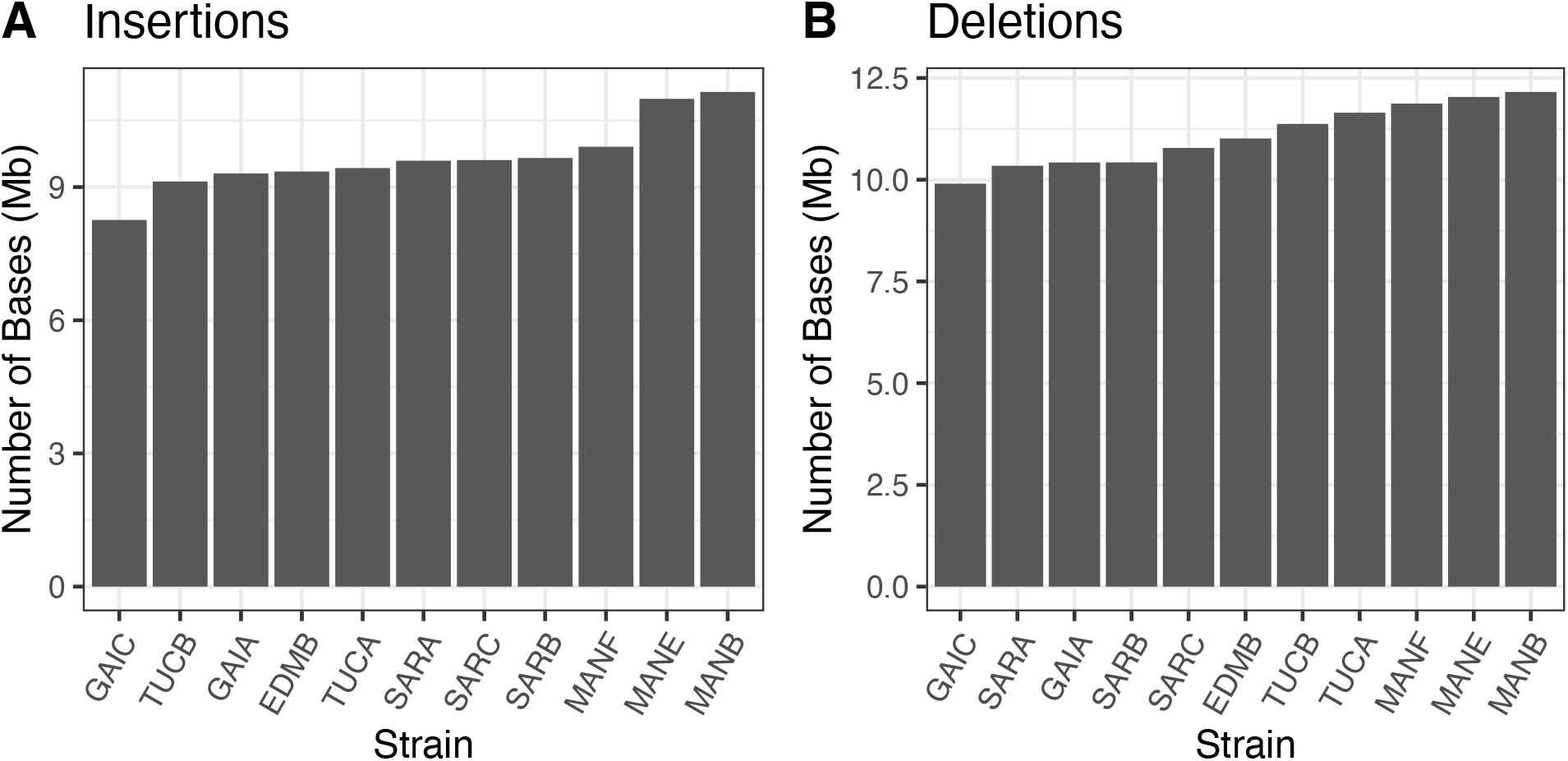
Numbers of bases impacted by (A) insertion and (B) deletion events in each Nachman strain relative to the mm39 reference sequence.

Many SVs in the Nachman strain genomes are potentially functional. Approximately 44% of SVs overlap RefSeq genes (23,733 SVs overlapping 10,051 unique genes), with 16,706 SVs overlapping annotated coding regions of 6368 unique genes. An additional 393 SVs are predicted to have high impact, with 214 ablating transcripts (**Supplementary Table 6**). The majority of these transcript ablating SVs impact predicted genes or pseudogenes, although several olfactory receptors, vomeronasal receptors, and immunoglobulins are harbor loss-of-function structural mutations.

Recent work has suggested that a majority of SVs in mouse genomes are due to transposable element activity (51). We theorized that many SVs in the Nachman strains are likewise mediated by TE-activity in the genome. Indeed, the size distribution of insertion and deletion calls in the Nachman strain call set is consistent with key contributions from TE activity (**Supplementary Figure 5**). To more formally assess this possibility, we annotated all insertion and deletion SV calls using repeatMasker. Overall, we identify 25.08 Mb of SV-associated sequence in the Nachman strains comprised of TE-derived sequence (13.82Mb deletion, 11.25Mb insertion), corresponding to 49.5% of SV-impacted bases. SINE B2 MM1A repeats are the most abundant TEs within SVs, (5612 polymorphic MM1A elements in the Nachman panel; **Supplementary Table 7**).

### Strain subspecies ancestry

An earlier investigation of wild-caught mice from Tucson, Arizona uncovered evidence for variable levels of introgression from *M. m. castaneus* (30). We used a sample of 29 wild-caught mouse genomes from *M. m castaneus*, *M. m. domesticus*, and the outgroup *M. spretus* to evaluate genomic evidence of possible *M. m. castaneus* introgression in the TUCA/NachJ and TUCB/NachJ lines (**Supplementary Table 8**). Patterson’s D is significantly non-zero for both strains (D_TUCA_ = 0.213 and D_TUCB_ = 0.192; P < 2.3 x 10^-16^ for both strains), with estimated *M. m. castaneus* admixture proportions (f_4_ ratio) of 13.4% and 11.3%, respectively. *D* is also significantly greater than zero for SARB/NachJ and SARC/NachJ (**Supplementary Table 9**). However, the estimated admixture proportion from *M. m. castaneus* is <1% in these strains, and may be attributable to the incomplete sampling of ancestral wild mouse diversity. We conclude that the genomes of both Tucson lines, but not strains from other locations, harbor significant *M. m. castaneus* ancestry. We also investigated possible introgression from *M. m. musculus* and found that all Nachman strains have <1% admixture from *M. m. musculus*, indicating no significant or recent introgression from this subspecies (**Supplementary Table 9**).

We next estimated admixture statistics in windows of 5000 informative SNPs (2500 slide) across the genomes of *M. m. castaneus* in TUCA/NachJ and TUCB/NachJ, focusing on the 5% of windows with the most extreme *f_d_* statistics to identify regions of likely *M. m. castaneus* introgression. We utilize *f_d_*, rather than the related *D* statistic, as the variance in *D* can be quite large when applied to small windows (53). Introgressed regions are overwhelmingly unique to either TUCA/NachJ and TUCB/NachJ (**Supplementary Tables 10 and 11; Supplementary Figure 6**), consistent with the high genomic divergence between these lines (**Figure 3**). For strain TUCA/NachJ, genes in introgressed regions are enriched for biological processes related to B-cell adhesion, alkaloid metabolism, lipid metabolism, sensory perception of bitter taste, regulation of systemic arterial blood pressure, and smell perception (**Supplementary Table 12**). For strain TUCB/NachJ, genes in regions of *M. m. castaneus* introgression are enriched for biological processes related to multiple aspects of immunity, cellular response to glucose starvation, AIM2 inflammasome complex assembly, response to lipopolysaccharide, and keratinization (**Supplementary Table 13**). Overall, introgressed regions are short, implying that introgression events are not recent (**Supplementary Figure 6**).

### Relationship to wild mouse diversity

To contextualize the variation present in the inbred Nachman strains with that observed in wild *M. m. domesticus*, we created a joint variant callset featuring the 11 inbred Nachman strains and 94 publicly available wild *M. m. domesticus* samples from multiple populations (Iran, France, Germany, and the Eastern United States). PC analysis on this Nachman-wild callset reveals genetic clustering of the Nachman lines with samples from Europe and the US, with samples from Iran isolated along PC1 (13.9% of the variance; **Figure 5A**). These trends are recapitulated by a maximum likelihood phylogenetic tree of these samples, which places the Iranian samples as ancestral to the Nachman strains and wild-caught mice from Europe and the United States (**Figure 5C**). Overall, our findings are consistent with previous population genetic analyses of wild mice, which suggest that *M. m. domesticus* from the Indo-Iranian valley harbor elevated genetic diversity (11,54). These findings also align with prior work indicating a European origin for North American house mice (55).

**Figure 5.**
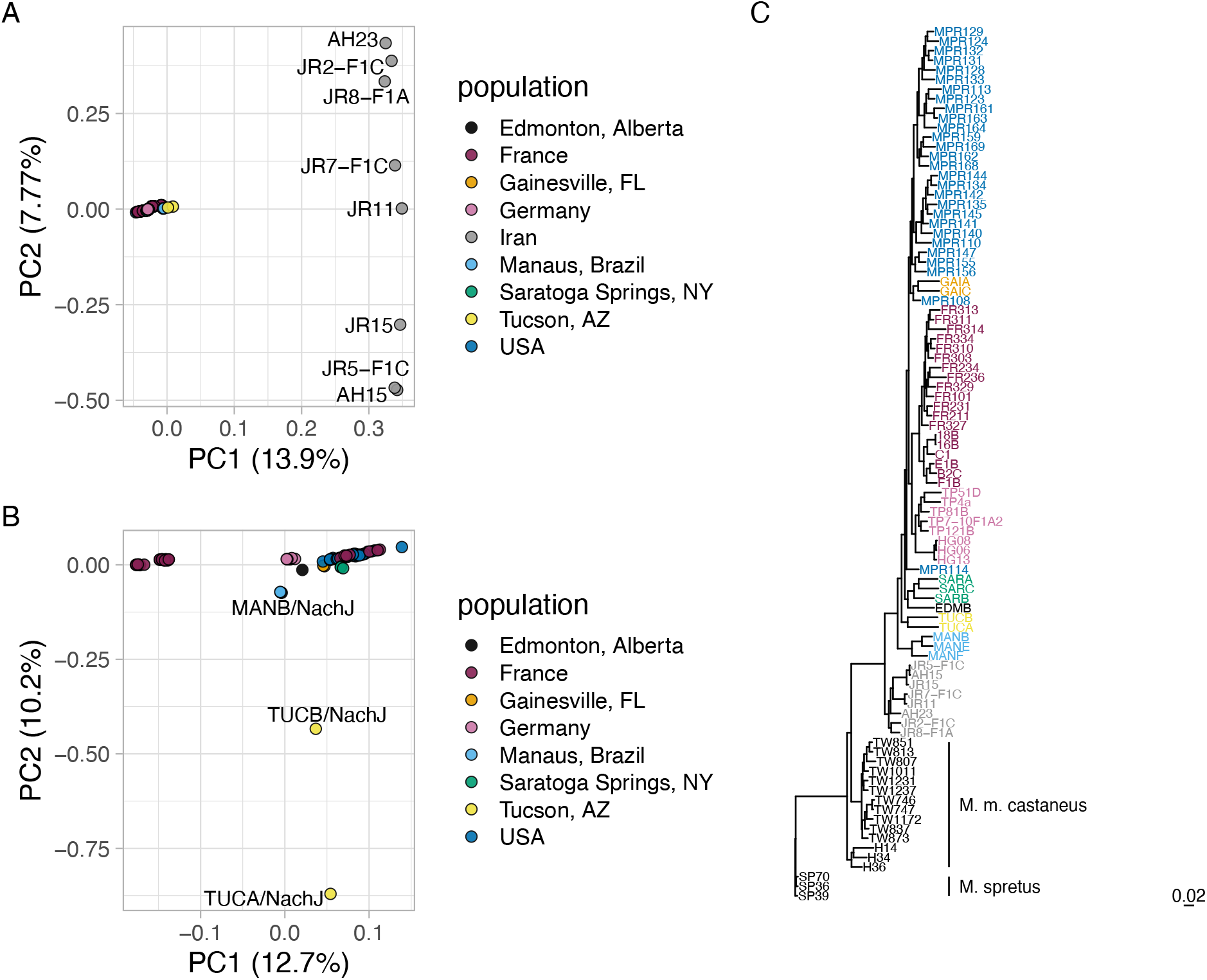
Nachman strains capture wild *M. m. domesticus* variation. (A) Principal component analysis performed on autosomal variants segregating in wild-caught *M. m. domesticus* mice from multiple populations and inbred Nachman strains. PC1 separates wild-caught mice from Iran and all other mice. PC2 stratifies mice from Iran. (B) Excluding mice from Iran provides increased granularity to detect differences across other *M. m. domesticus* populations. PC2 isolates the Nachman strains from Tucson, Arizona from other strains and populations. (C) Maximum likelihood tree constructed from biallelic chr19 SNPs. Strains are color-coded according to the legends in A and B.

We excluded the Iranian samples and repeated the PCA to evaluate how the genetic diversity captured in the Nachman samples compares to the genetic variation in contemporary *M. m. domesticus* from Europe and the Eastern US (**Figure 5B**). PC1 (12.7% of the variance) stratifies mice from wild-caught *M. m. domesticus* populations, with Nachman lines falling at intermediate positions along this dimension. PC2 (10.2% of the variance) isolates the two Nachman strains from Tucson, likely reflecting the presence of *M. m. castaneus* admixture in these lines. While the Nachman lines introduce significant new variation into laboratory strain collections, the 11 inbred strains in this panel sample only a subset of the genetic variation present in wild *M. m. domesticus* populations.

### Nachman wild-derived inbred strains capture extensive phenotypic diversity

We subjected mice from the majority of JAX imported Nachman lines to a 19-week phenotyping pipeline to profile strain variation in multiple metabolic, neurobehavioral, physiological, morphological and biochemical traits (**Figure 6**). Males and females from 9 strains were phenotyped across 16 cohorts of age-matched animals (+/-6 days). Cohorts were comprised of mice from multiple strains, and the majority of phenotyping cohorts included C57BL/6J control mice to permit post-hoc detection of potential batch effects. The strain composition of each cohort is provided in **Supplementary Table 14**. Overall, we collected 1119 phenotype measures from each animal, although many trait values are highly correlated (**Supplementary Table 15**).

**Figure 6.**
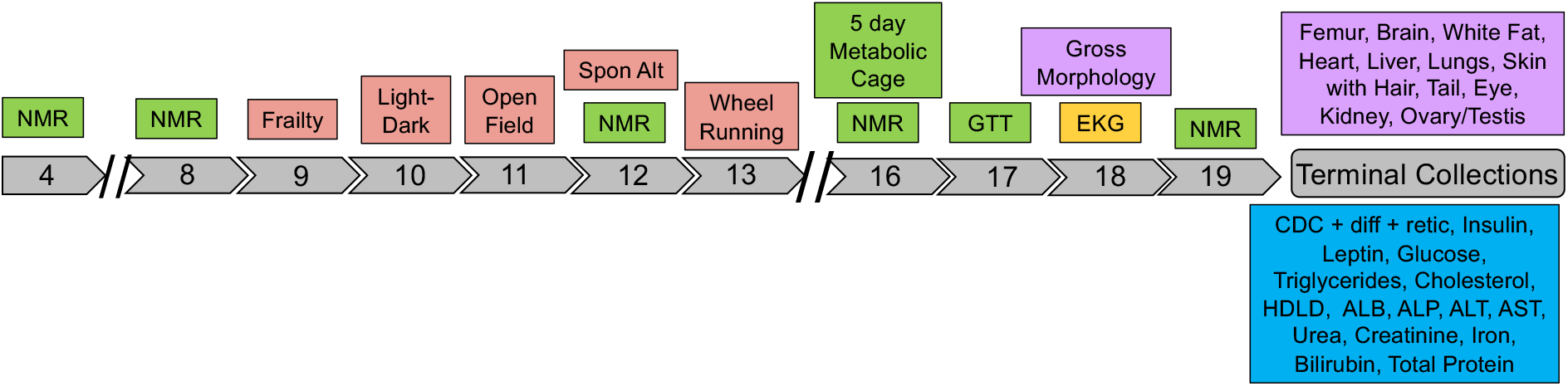
Schematic of the phenotyping pipeline. Mice were transferred to the Center for Biometric Analysis (CBA) at JAX at 4 weeks of age. Nuclear magnetic resonance (NMR) was used to assess body composition at 4, 8, 12, 16, and 19 weeks. From weeks 9-13, animals were subject to a series of neuro-behavioral testing paradigms, including light-dark test, open field, spontaneous alternation with Y-maze, and voluntary wheel running. From weeks 16-17, animals were subject to 5-day indirect calorimetry trials and intraperitoneal glucose tolerance testing. At week 18, mice underwent an unconscious EKG to assess cardiac rhythm and function. Mice exited the phenotyping pipeline at 19 weeks. Harvested blood and plasma samples were used to assess multiple biochemical traits and tissues were dissected. A subset of tissues were frozen for future molecular analysis and others were paraffin embedded for future histological study.

More than 90% of the 1119 surveyed phenotypes differ among Nachman strains, with 85.2% and 78.8% of phenotypes differing significantly in comparisons involving only females or males, respectively (Kruskal-Wallis test, P<0.05; **Supplementary Table 16**). We obtain qualitatively identical results using both one-way ANOVA (**Supplementary Table 17**) and a linear mixed effect modelling approach with strain treated as a random factor (90.9% of models including strain as a mixed effect variable provide significantly better model fit than a reduced model excluding strain; see Methods; **Supplementary Tables 18 and 19**). Approximately 25% (n = 290) of phenotypes exhibit significant differences between males and females and a significant strain-by-sex effect is observed for 21% (n = 237) of surveyed measures (two-way fixed effects ANOVA, P < 0.05; **Supplementary Table 20**). Similar trends are again observed using a linear mixed effects model comparison strategy: 37.0% and 20.5% of models including sex and interaction terms, respectively, provide improved fit compared to simpler models that exclude these effects (**Supplementary Tables 18 and 19**). Thus, phenotypic variation is ubiquitous across the Nachman strain panel, and many traits vary between sexes in a strain-dependent manner.

Below, we highlight results from each phenotype testing paradigm, present estimates of broad sense heritability, and compare the phenotypic variance among the Nachman samples to that present across classical inbred mouse strain panels. We then present results from correlation analyses that cut across multiple phenotype domains. All phenotype data will be deposited on the Mouse Phenome Database and are available in **Supplementary Table 21**. Strain-level and Strain x Sex-level phenotype means are provided in **Supplementary Table 22**. Broad-sense heritability estimates derived from one-way ANOVA (Phenotype ∼ Strain) for each phenotype are presented in **Supplementary Table 17**. Results from non-parametric Kruskal-Wallis tests of strain effects on each phenotype are provided in **Supplementary Table 16**. Two-way ANOVA results (Phenotype ∼ Strain * Sex) are presented in **Supplementary Table 20**. Results from linear mixed effects model fitting are provided in **Supplementary Tables 18 and 19**.

#### Body composition analysis by NMR

We assessed multiple metrics of body composition (body weight, fat mass, lean mass, water mass) via NMR at 5 timepoints between 4 and 19 weeks of age. As expected, all mice gained weight during this period, although the percentage increase in body mass differed among strains across this 15 week interval (range: 28.1% SARC males to 95.92% in GAIC/NachJ males). Males trended toward larger body mass than females (Average sex dimorphism across strains: 4.92 g; **Figure 7**), and all Nachman lines are smaller than C57BL/6J control mice. The broad sense heritability of body weight at 19 weeks is 0.89 in both males and females (**Supplementary Table 17**), indicating that significant proportion of strain variation in body mass is attributable to underlying genetic differences between strains.

**Figure 7.**
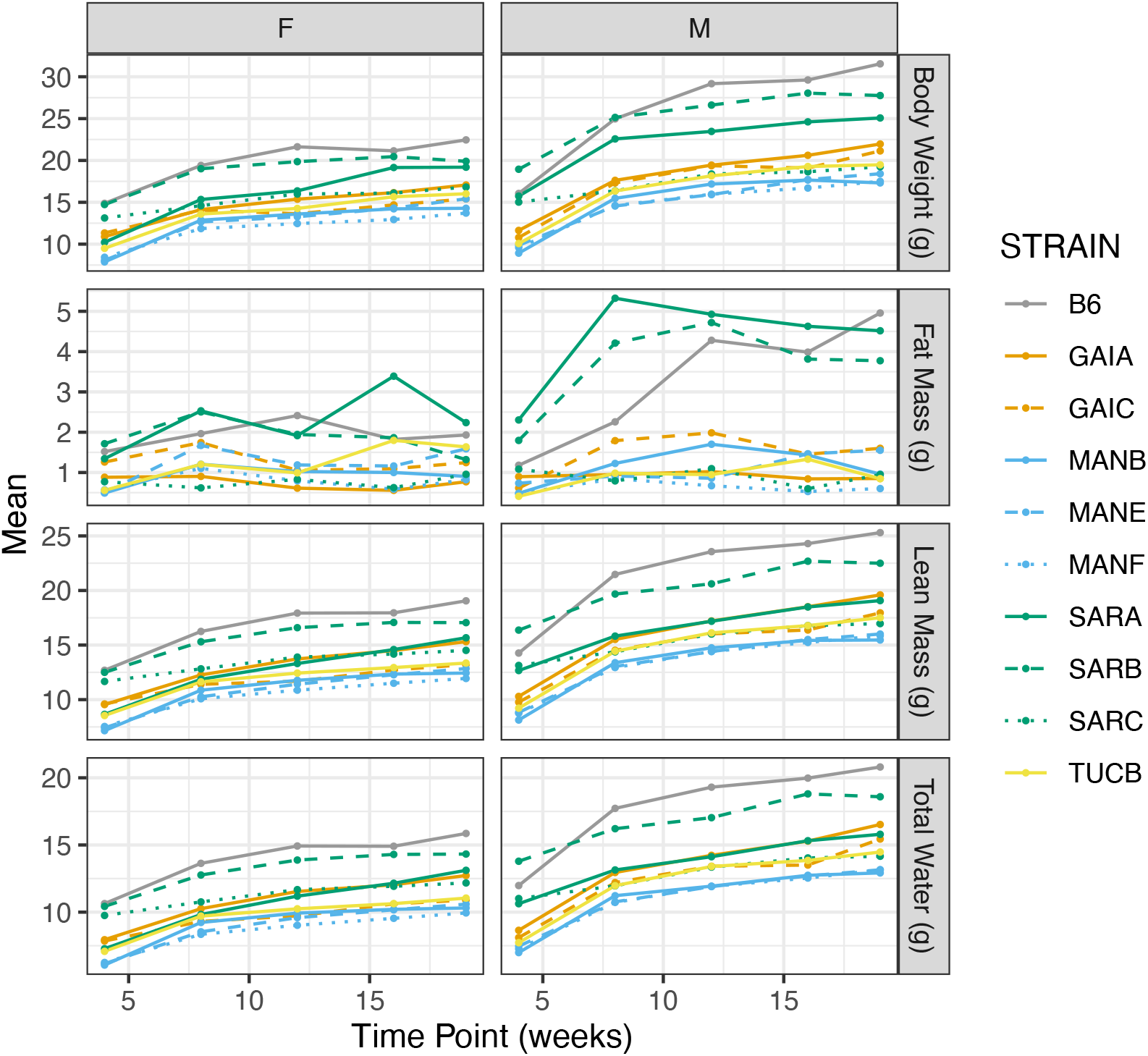
Changes in body composition over time as assessed by NMR. Points correspond to strain-level means and data are partitioned by sex to highlight the degree of sex-dimorphism in body composition. Strains are color coded by geographic origin, with strains from a common location distinguished by line type.

We similarly document significant strain and sex differences in body fat mass, lean mass, and water mass (**Supplementary Tables 16,17,20**; **Figure 7**). Overall, body composition phenotypes are highly correlated over time within strains (average Spearman’s rho = 0.67; **Supplementary Data 23**), implying that increased body mass is driven by coordinated changes in fat, lean, and water mass rather than isolated changes in one compositional component. Fat mass is the most volatile compositional component, with several strains exhibiting negative correlations over time (**Supplementary Table 23**). Body composition measures are correlated across strains at individual timepoints (all Spearman’s rho > 0.3), consistent with the observation of near parallel growth trajectories across strains (**Supplementary Figure 7**).

#### Frailty Assessment of Overall Body Condition

Overall health and body condition were assessed using a 29-dimension frailty index score (56). All strains have low mean frailty index scores (values ranging from 1.75-3.0 out of a possible score range of 0-28; **Supplementary Table 22**), although the qualitatively small strain differences observed exceed chance expectations (Kruskal-Wallis *P* = 0.01237; **Supplementary Table 16**). Body temperature differed significantly across strains, with GAIC/NachJ and the three strains from Saratoga Springs, NY exhibiting the highest values (Kruskal-Wallis; *P* =2.39 x 10^-6^; **Supplementary Tables 16 and 22**).

#### Light-Dark test for anxiety-like behaviors

The light-dark test is premised on the natural aversion of rodents to being in brightly lit spaces; the proportion of time mice spend in the brightly lit versus dark zones of the testing chamber yields quantifiable metrics of anxiety-like behaviors. Strain TUCB/NachJ shows the lowest overall ambulatory time (24.5 ± 3.54 s; range of other strain means: 30.2-35.8 s), fewest crossovers from one zone to the other (30.9 ± 12.2; range for other strain means: 31.8-60.6), and the greatest proportion of time in the dark chamber zone (67.1%; range for other strain means: 48.0-59.5%; **Supplementary Table 22**), consistent with higher overall levels of anxiety-like behavior in this strain. MANE/NachJ spent the greatest proportion of time in the light zone (52%), whereas GAIA/NachJ traveled the greatest total distance in the light zone (819 ± 159 cm; range of other strain means: 436 – 745 cm). Overall, females are more active in the dark than males (Total Distance Traveled in Dark, F_1_ = 7.88, P = 0.0055), with an especially pronounced sex dimorphism for a number of anxiety-like traits in SARA/NachJ (**Supplementary Figure 5**). Importantly, the time spent at rest and number of bouts of movement in the light are moderately heritable (*H^2^* > 0.3; **Supplementary Table 16**), revealing a genetic component to observed strain variation.

#### Open Field Assay to quantify overall locomotion and exploratory behavior

Most measures of locomotion and exploration exhibit high broad sense heritability in the Nachman lines (**Supplementary Table 17**). Strains vary more than 3-fold in total distance traveled within the arena over a 60 minute test period, ranging from a low of 5158 ± 803 cm in GAIC/NachJ to a high of 17141 ± 4135 cm in SARC/NachJ (F_9_ = 24.9; *P* = 1.46×10^-27^). We observe significant sex effects on several open field metrics, including the number of independent movement episodes and vertical activity time (**Supplementary Table 20**). Females exhibited more bouts of movement (Females: 634 ± 101 episodes; males 594 ± 118 movement episodes; F_1_ = 11.24; *P* = 0.00097), whereas males spent more time moving in the vertical plane (males: 330 ± 194 s; females: 274 ± 150 s; F_1_ = 7.16; *P* = 0.0081). We report a significant interaction between sex and strain for the total time distance traveled in the center of the arena (F_9_ = 2.099; P = 0.032), with SARA/NachJ and SARC/NachJ females traversing significantly greater distances than their male counterparts (Wilcoxon Rank Sum Exact Test; *P* < 0.05; **Supplementary Table 20**).

#### Spontaneous Alternation in the Y Maze

The spontaneous alternation assay is premised on the natural tendency of mice to explore novel environments and serves as a test of spatial working memory. Nachman strains vary in the total number of arm entries and number of spontaneous alternations in the Y-maze, with both phenotypes also exhibiting significant sex and strain x sex effects (Two-way ANOVA, *P* < 0.05; **Supplementary Table 20**). In particular, SARA/NachJ mice show the greatest number of spontaneous alternations (48.33 ± 7.37; range for other strains: 23.85-34.80), with females exhibiting higher numbers than males (SARA/NachJ males: 42.13). Although this difference is not significant owing to the modest number of mice tested (n = 3 SARA/NachJ females; **Supplementary Table 14**), our finding is concordant with the discovery of increased exploratory behavior in SARA/NachJ females in the open field assay. The alternation ratio (percent alternation) for most strains exceeds that for C57BL/6J, suggesting that the Nachman lines have stronger working memories and/or a more intense innate desire to explore novel spaces than C57BL/6J.

#### Wheel Running

At week 13, mice were subjected to a 3-day home cage wheel running assay to assess overall activity and circadian rhythms. Strains vary 6.8-fold in total distance run over the 60h trial, from a low of 9,112 m in MANE/NachJ to 62,019 m in SARC/NachJ (**Supplementary Table 22**). Mice are nocturnal and, as expected, activity was highest during nighttime hours for all strains (Wilcoxon Rank Sum Test, P < 0.0005; **Figure 8A**).

**Figure 8.**
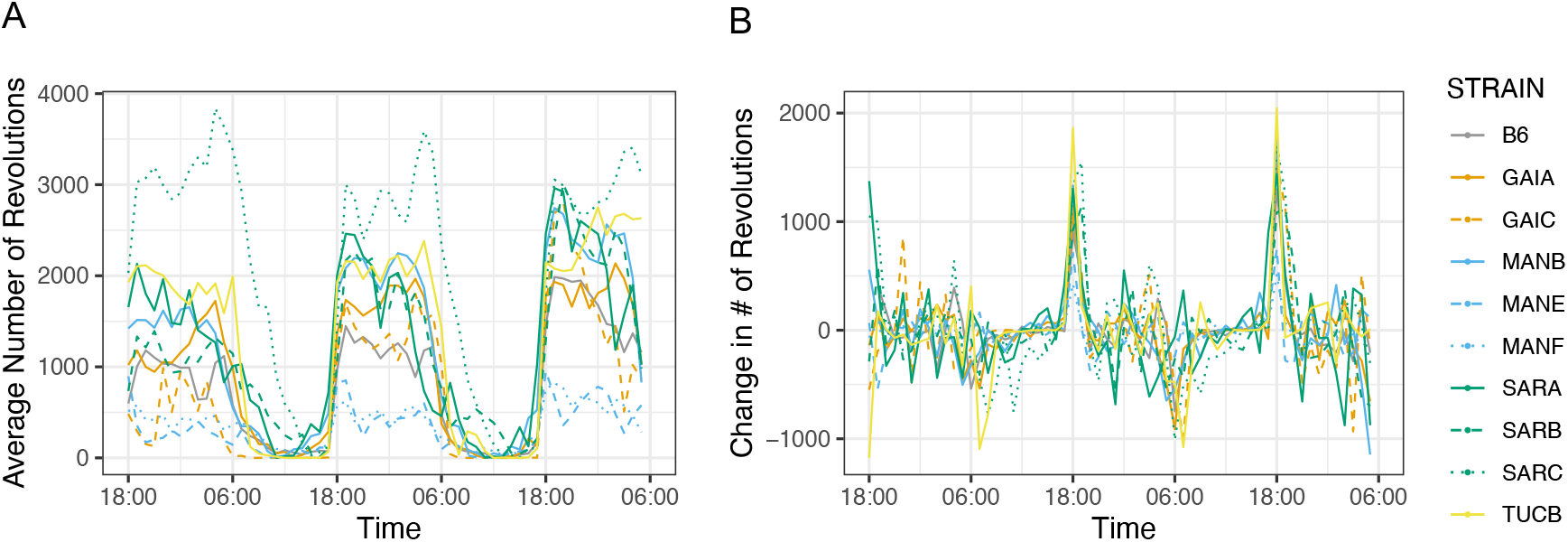
Strain variation in voluntary exercise and circadian rhythm. (A) The average number of wheel revolutions per strain over a 60 hour trial period. (B) Hour-to-hour change in wheel running activity (number of wheel revolutions). Abrupt changes in activity correspond to transitions between sleep-wake cycles. In both figures, strains are color-coded by geographic origin, with strains from a common locations further distinguished by line type.

The Nachman strains derive from wild-caught mice exposed to distinct photoperiods in the wild. We sought to determine whether strains differ in the timing of their peak wheel running as a function of geographic origin. To this end, we computed the change in the number of wheel revolutions per hour across the full trial, defining the transition between day and night as the point exhibiting the most extreme difference in activity level. All strains commence nighttime levels of activity at ∼18:00h, but the onset of daytime rest periods is considerably more variable, with strain differences in the duration of nighttime exercise and daytime activity. For example, mice from Manaus, Brazil exhibit a shorter duration and reduced levels of nighttime activity compared to strains from Saratoga Springs, New York (**Figure 8B**), which remain active during much of the day.

#### Indirect calorimetry

Mice were subject to continuous, high-definition respiratory monitoring in Promethion Core cages for 5 days, allowing estimation of energy expenditure, activity levels, and food consumption. We observe significant strain effects on all surveyed phenotypes (P < 0.0001), with approximately half of the metabolic traits captured in this assay also exhibiting significant differences between the sexes (**Supplementary Tables 16, 17, 20**). Both sexes of all strains were most active, exhibited highest energy expenditure, and consumed the most food during the dark cycle (**Figure 9A-C**; Paired Wilcoxon Signed Rank Test, all *P* < 10^-10^). Most metabolic traits are highly heritable (**Supplementary Table 17**), nominating the Nachman lines as excellent models for genetic studies of metabolism. Intriguingly, despite their close genetic proximity, the two strains from Gainesville, Florida exhibited the lowest (GAIC/NachJ; 7.53 kcal/24h) and highest (GAIA/NachJ; 9.98 kcal/24h) total energy expenditure, with C57BL/6J controls presenting an intermediate phenotype (9.67 kcal/24h (**Figure 9D**).

**Figure 9.**
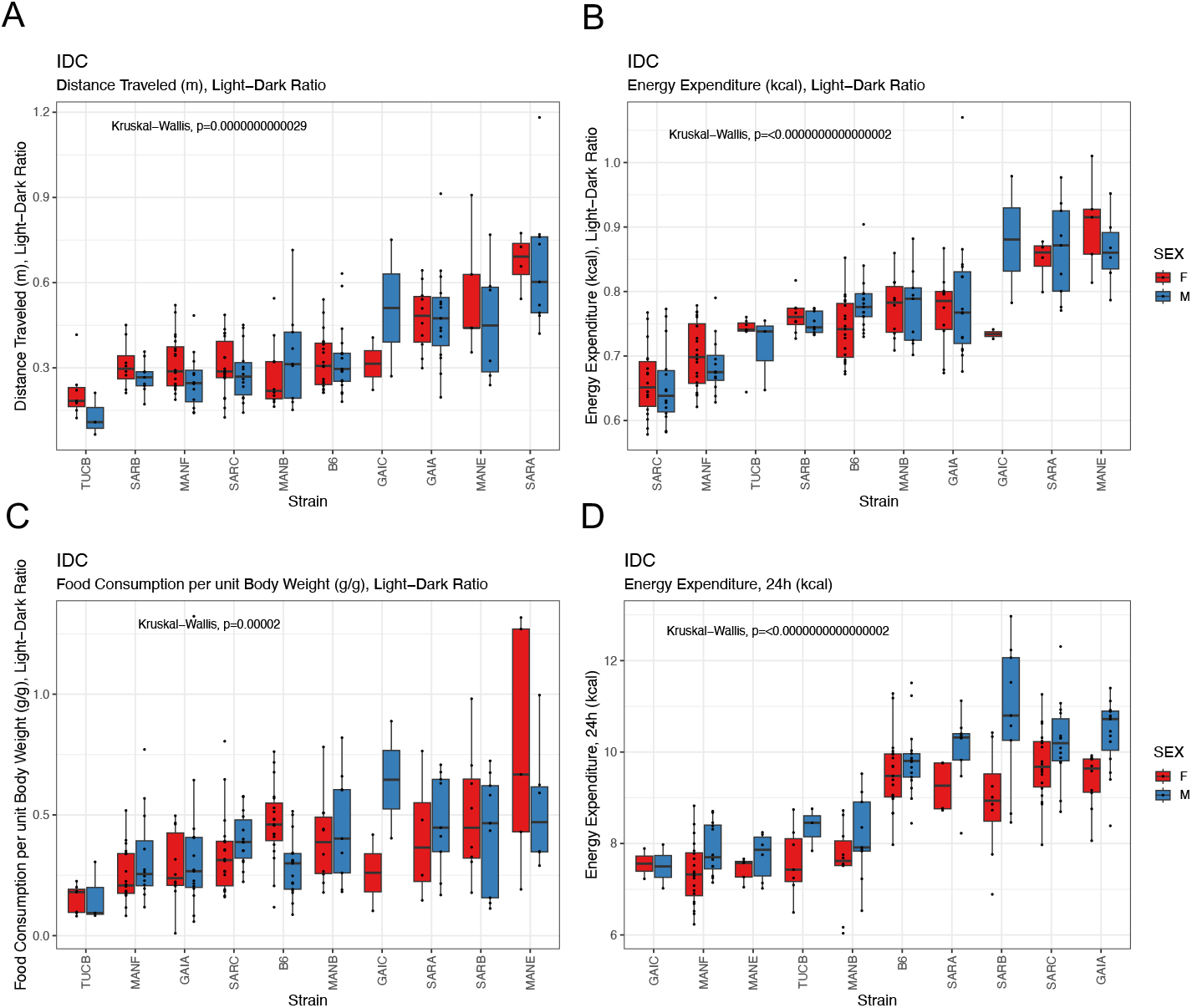
Strain differences in metabolic phenotypes quantified by continuous respiratory monitoring. (A) All strains exhibit higher activity during the night, although the exent of the nighttime activity bias varies across strains. (B) Similarly, all strains expend more energy and (C) consume more food at night, but the ratio of day:night energy expenditure and food intake varies across strains. (D) Total energy expenditure varies significantly across strains, with the two strains from Gainesville, Florida (GAIC/NachJ and GAIA/NachJ) delimiting the extremes.

#### Glucose Tolerance Test

All Nachman lines show an attenuated response to intraperitoneal injection of a controlled concentration of glucose relative to C57BL/6J (**Figure 10**). Peak blood glucose concentrations are higher in males than females for all strains, with exceptionally pronounced dimorphisms in SARB/NachJ and GAIC/NachJ (range of sex dimorphism across strains at 15 minutes post injection: 5.29-91 mg/dL). Glucose response curves are significantly different for all pairwise strain comparisons involving SARC/NachJ males and females (Permutation test *P* < 0.005, **Supplementary Table 24; Figure 10**), emphasizing the unique response profile of this strain. The area under the glucose response curve varies 2.4-fold across strains, with strains GAIC/NachJ and C57BL/6J delimiting the lower and upper extremes, respectively (**Supplementary Table 22**).

**Figure 10.**
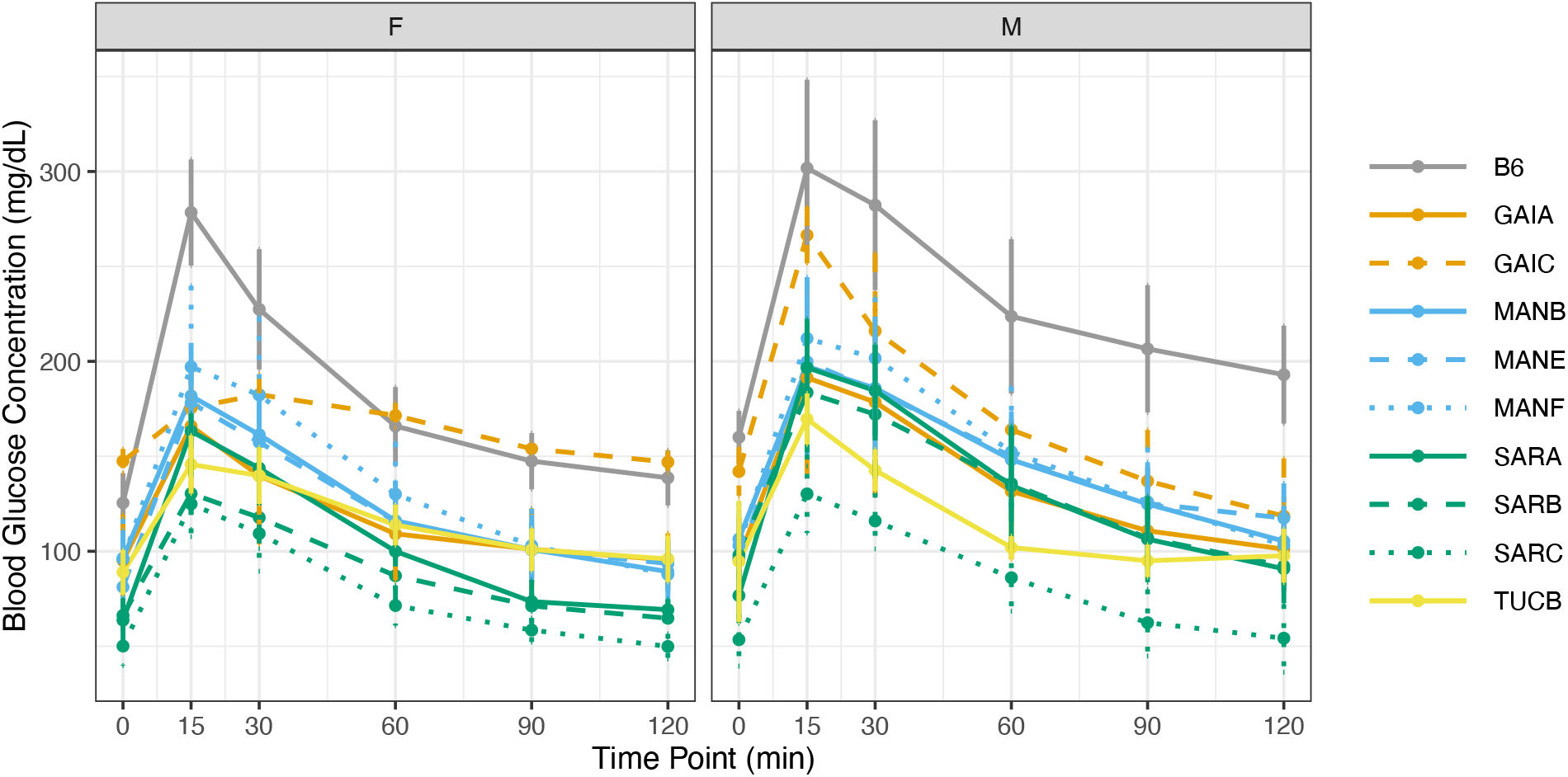
Glucose response curves over a 2-hour window. Males and females were analyzed separately. Mice were fasted for 4 hours prior to administration of a fixed (w/v) amount of glucose by intraperitoneal injection. Blood glucose levels were assessed at 15, 30, 60, 90, and 120 minutes post injection. In both panels, strains are color-coded by geographic origin, with strains from a common location further distinguished by line type. Vertical bars denote +/-1 SD around the strain mean.

#### Unconscious Electrocardiogram

Mice underwent unconscious electrocardiogram (EKGs) at week 18 of the phenotyping pipeline (**Figure 6**) to assess heart rate and rhythm. Strains exhibit significant variation in all surveyed EKG metrics, including heart rate, peak amplitudes, wave durations, and peak intervals (**Supplementary Tables 16**; Kruskal-Wallis *P* < 0.005). Nachman lines have lower heart rates, smaller PR intervals, and Q amplitudes than C57BL/6J control animals (**Supplementary Table 22**), suggesting clinically relevant differences in cardiac function between laboratory and wild mice.

#### Gross Morphology and Organ Weights

Nachman strains exhibit significant, heritable variation in body length and tail length (Kruskal-Wallis 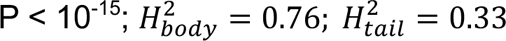; **Supplementary Tables 16 and 17**; **Supplementary Figure 6**). Both phenotypes exhibit a significant sex effect, with males having both longer bodies and longer tails than females (**Supplementary Table 20)**. The magnitude of the sex dimorphism in both morphology traits varies by strain, with SARA/NachJ and SARB/NachJ males exhibiting much longer body lengths than their female conspecifics and only a modest length dimorphism between MANB/NachJ males and females (**Supplementary Figure 9**). Similarly, we observe significant heritable variation in organ weights among strains and between the sexes (**Supplementary Table 17**). Brain size exhibits exceptionally high heritability (H^2^ = 0.75) and variability among strains, even after standardizing by total body weight (Kruskal-Wallis P = 6.2×10^-11^), revealing that observed brain size differences among strains are not simply proportional to overall body size.

#### Clinical Chemistry

We assessed levels of numerous blood-based clinical markers after subjecting mice to a 4hr fast at 19 weeks (**Figure 6**). Nachman strains vary in immune cell populations, blood lipid profiles, measures of liver function, and both platelet and red blood cell composition (**Supplementary Tables 16 and 17**). For example, total cholesterol levels range 2.7-fold among strains, with TUCB/NachJ and GAIC/NachJ mice defining the extremes (71.2 mg/dL – 195 mg/dL). Males from all strains have higher total cholesterol values than their female conspecifics (average dimorphism = 35.8 mg/dL), a trend echoed in levels of HDLD cholesterol (**Figure 11A,B**). Nachman lines show exceptionally high variability in platelet counts (range of strain means: 754-1725 x10^3^ cells/uL) and platelet volume (range: 6.39-9.58 fL), with lines from Manaus, Brazil presenting high numbers of low volume platelets, and strains from Saratoga Springs having low numbers of high volume platelets (**Figure 11C-D**). Multiple immune cell populations, including monocytes, lymphocytes, neutrophils, and eosinophils, exhibit heritable strain variation across the Nachman strain panel (**Supplementary Table 17**), consistent with recent reports of strain differences in pathogen response (57).

**Figure 11.**
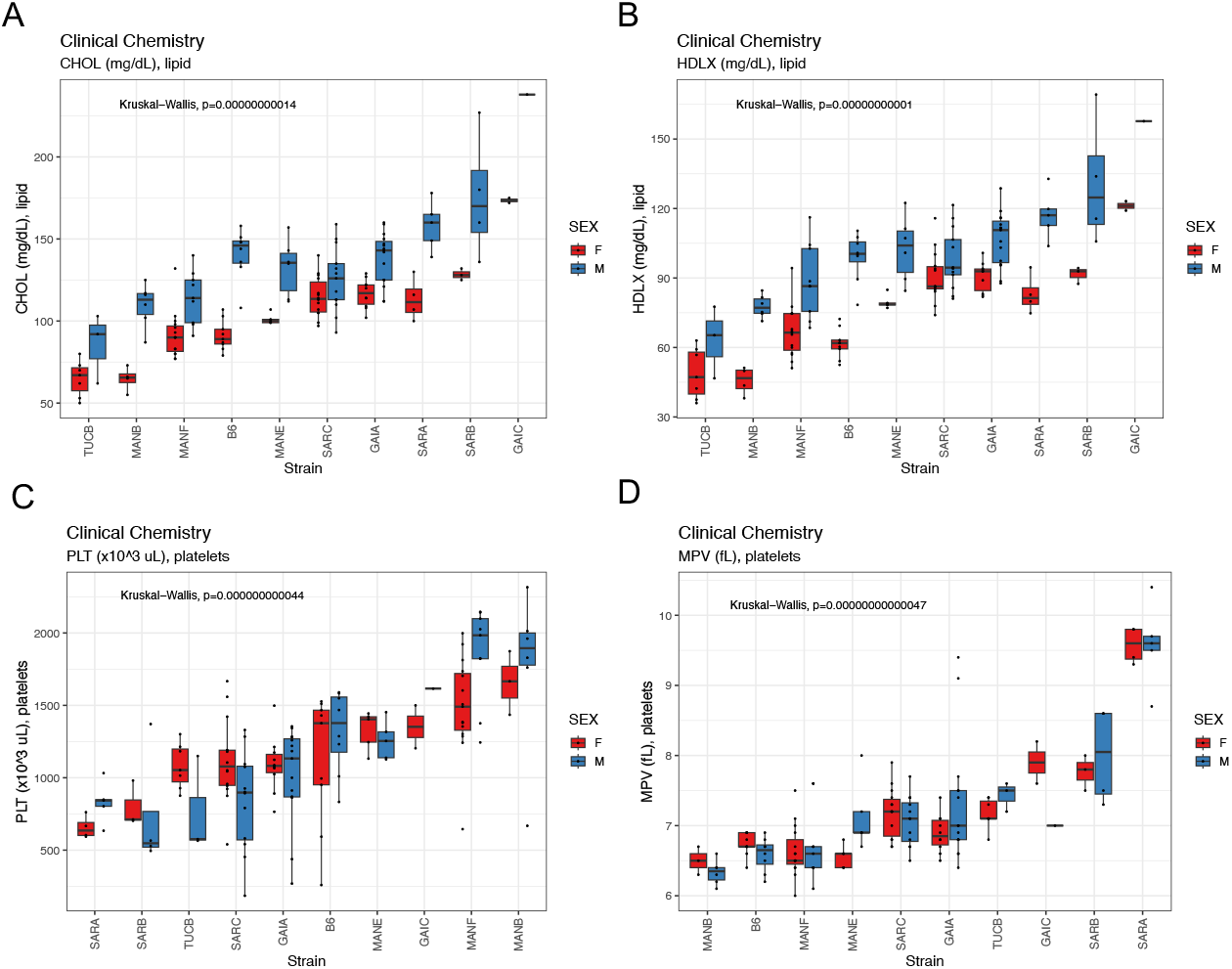
Boxplots depicting the extent of strain and sex variation in (A) total cholesterol, (B) HDLD cholesterol, (C) platelet count, and (D) platelet volume.

### Integrated analysis of Nachman and classical inbred strain phenotypes expands the variance observed in classical inbred strains alone

We intersected phenotype data from the Nachman lines with existing strain survey datasets deposited on the Mouse Phenome database (see Methods). The Nachman animals expand the range of phenotype values observed across inbred strain panels for several phenotypes, including total platelet counts, percent fat mass, and hematocrit levels (**Supplementary Figure 10**).

The inclusion of C57BL/6J mice as controls in our phenotyping cohorts provides a common touchpoint between our data and previously published datasets. While we find that C57BL/6J trait values are generally stable across phenotyped cohorts (**Supplementary Table 25**), C57BL/6J trait values differ significantly between datasets (**Supplementary Tables 26 and 27**). The inconsistency in trait values could owe to differences in animal age at the time of phenotyping, differences in housing conditions, animal handling by different technicians and research staff, differences in phenotyping protocols or equipment, or other vagaries of the experimental environment. Regardless of the source of this phenotypic variance, it poses major obstacle to phenotype data integration with the Nachman strains and underscores the need for caution in the interpretation of absolute differences in trait values across independently collected datasets.

### Strain-level trait correlations across phenotype assays

Significant strain-level phenotype correlations may reveal traits regulated through shared genetic pathways. As expected, measures of body composition, body size, and organ weights are positively correlated, suggesting a general pattern of isometric growth (**Supplementary Figure 7**). Additionally, measures of total activity in indirect calorimetry and open field assays are positively correlated (**Supplementary Table 15**), as expected. Total serum bilirubin levels are negatively correlated with several phenotypes assessed from the spontaneous alternation assay, including total number of arm entries (rho = −0.95; P = 0) and the number of spontaneous alternations (rho = −0.78, P = 0.01165), but positively correlated with the alternation ratio (rho = 0.65; P = 0.049). This latter result confirms published reports of a negative association between serum bilirubin concentration and cognition impairment in schizophrenia in humans (58), extending this correlation to mice. Leptin levels are positively correlated with fat mass at each timepoint assessed by NMR (4, 8, 12, 16, and 19 weeks; rho > 0.74; P < 0.018) and gonadal fat pad weight (rho = 0.68; P = 0.035), and are negatively correlated with food consumption (rho = - 0.87; P = 0.0027). These results reinforce the well-established link between leptin secretion and satiety sensing. The duration of the QRS interval on an unconscious EKG is positively correlated with food consumption, fat mass, and body size across strains. This result lends independent support to a recently published association between QRS duration and BMI in humans with no overt cardiac disease (59). A comprehensive summary of all pairwise trait correlations is provided in **Supplementary Table 15**. Sex-specific trait correlations are provided in **Supplementary Tables 28 and 29**.

## DISCUSSION

Here, we introduce a new inbred mouse strain resource, the Nachman wild-derived inbred strain panel. Strains in this panel derive from wild-caught mice adapted to unique environments across North and South America. Integrated analysis of the genomes of these new strains with those of the classical inbred strains reveals millions of novel genetic variants, including predicted deleterious alleles and gene-spanning structural variants. Paralleling this genetic diversity, Nachman strains capture considerable phenotypic variation across biochemical, neurobehavioral, physiological, morphological, and metabolic trait domains.

We attempted to integrate our phenotype data with publicly available phenotype datasets from prior mouse strain surveys. Importantly, the C57BL/6J controls included in our phenotyping cohorts provide a common data point to anchor our data to published phenotyping efforts. However, due to differences in animal ages, experimental methodology, and environmental housing conditions, the vast majority of phenotypes present inconsistent trait measurements across C57BL/6J, limiting the extent to which we can integrate data from the Nachman lines with other strain panels. These challenges underscore well-known difficulties with data integration and emphasize the importance of standardized phenotyping protocols and detailed metadata reporting (60). While we acknowledge these caveats, our findings place high certainty that the Nachman strains extend the range of trait values realized using classical inbred strains alone (**Supplementary Table 10**).

The Nachman strain panel complements existing diverse mouse populations, such as the BXD, CC, HGMP, and DO, offering a new community resource for profiling phenotypic variation and assessing responses to exposures, interventions, or treatments. While the 11-member panel is too small to power genetic mapping studies, experimental crosses between Nachman strains with extreme phenotypes can be employed to generate F2, backcross, or advanced intercross lines for experimental mapping.

At the same time, a key feature that distinguishes the Nachman panel from other mouse diversity resources is its exclusive profile of natural genetic and phenotypic variability in *M. musculus*. During the creation of these wild-derived strains, attention was paid to minimize the impacts of artificial selection, thereby assuring that the resultant fixed strain genomes most faithfully mirror haplotypes found in the wild. In contrast, laboratory inbred strains are products of strong selection for morphological traits, increased reproductive output, and behavioral traits that promote ease of handling. Such strong artificial selection for traits of interest can have broad genomic consequences on unrelated phenotypes through pleiotropy and linkage (61). The deliberate efforts undertaken during the generation of the Nachman lines help to assure that these strain genomes immortalize multiallelic patterns of diversity that most closely approximate the polygenic architecture of traits in nature, including the complex genetic basis of human disease-related phenotypes.

Our phenotyping efforts recapitulate adaptive phenotypic differences documented in wild house mice. For example, we find that SAR mice are typically bigger, more active, and have higher energy expenditure than MAN mice, consistent adaptative phenotypic evolution to colder environments (32) and predictions of Bergmann’s Rule (62). Importantly, the impact of local adaptive pressures on phenotype diversity across the Nachman strains reveals strong parallels to humans and enhances the potential promise of this resource for robustly modeling human phenotypes. Notably, selection is a crucial driver of phenotypic divergence between human populations, including differences in disease susceptibility and resistance (63).

Analyses of subspecies affiliation and introgression reveal that strains from four of the five sample locations (Manaus, Brazil; Saratoga Springs, New York; Gainesville, Florida; and Edmonton, Alberta, Canada) are of pure *M. m. domesticus* ancestry, but point to moderate levels of introgression from *M. m. castaneus* in mice from Tucson. This finding adds to prior observations of *M. m. castaneus* introgression into mouse populations from the West coast (64) and earlier SNP-based genetic investigation of mice from the American southwest (65). Importantly, there are no known hybrid incompatibilities between *M. m. castaneus* and *M. m. domesticus*, and our early assessments of F1 hybrids from crosses involving the Tucson lines and other Nachman strains reveals no overt fertility deficits (**Supplementary Table 3**). Thus, we consider it unlikely that infertility or inviability will present challenges to genetic crosses between Nachman strains, between Nachman strains and classical inbred strains, or between Nachman strains and other wild-derived inbred strains of *M. m. domesticus* or *M. m. castaneus*.

Our initial phenotypic and genetic characterization of these new strains provides the basis for many new discoveries and establishes the foundation for future efforts to build upon. For one, all phenotyping was performed “at baseline” using animals fed a standard rodent diet, housed under standard conditions, and in the absence of any experimental treatments. Experimental designs that employ environmental perturbations, different exposure or treatment regimes, aged animals, or other deliberate manipulations may uncover novel phenotypic responses or resilience/susceptibility traits in the Nachman strain panel. Second, our phenotyping pipeline was purposefully designed to survey a broad range of biomedically relevant traits and is far from comprehensive. For example, surveyed phenotypes exclude sensory perception phenotypes, bone density, histopathology assessments, and microbiome composition. At the same time, deeper, more exhaustive phenotyping of specific trait domains could unlock strain-level differences that fail to manifest using coarser phenotyping assays. Third, due to COVID-related impacts, mice from several strains were poorly represented (GAIC/NachJ, TUCA/NachJ, TUCB/Nach) or altogether absent (EDMB/NachJ) from our phenotyped cohorts (**Supplementary Table 14**). Thus, the extent of phenotypic variation across Nachman strains presented here is potentially underestimated – a possibility that future investigations could assess. Fourth, we sequenced each strain to modest coverage and relied on comparisons to the mm39 reference genome for variant discovery. We obtained only low-coverage of the X and Y chromosomes in our sequenced males (∼5x), requiring us to exclude these chromosomes from our genomic analyses. Ultimately, we endeavor to perform additional long- and ultra-long read sequencing, generate high quality de novo sequence assemblies, and perform assembly-based variant calling to more comprehensively catalog variation in these strain genomes.

We also acknowledge practical caveats to the use of mice in this strain panel. Our colony breeding records demonstrate that the Nachman strains are robust breeders, but breeding performance falls short of what is observed for many classical inbred strains that have been subject to decades of artificial selection for productivity (**Figure 2**). Further, like many other wild-derived inbred strains, breeder dams from the Nachman lines do not possess externally visible mating plugs, potentially due to their rapid loss or re-absorption following mating. This may pose challenges for experiments that hinge on the ability to precisely time matings or obtain fetuses at specific gestational ages. Males from many strains also exhibit high levels of aggression toward cage mates and may require single housing or additional enrichment to minimize aggressive encounters. Anecdotally, we have found that supplying mice with wooden blocks for gnawing minimizes tail biting among cage mates. Finally, strains in the Nachman panel retain very high levels of wildness, imposing challenges to routine handling. Of special note, juvenile mice from the TUCB/NachJ strain are remarkable jumpers, easily capable of escaping cages topped with cage extenders (total height ∼14”). While working with wild mice may be daunting at first, it is our experience that competency and confidence build quickly. Specialized workstations can aid in containing mice during routine handling, and innovations in animal housing could eventually allow for hands-off cage changes (e.g., via attachment of open plastic tubing to closeable ports on dirty and clean cages, allowing mice to move into new cages on their own). Further, continuous home-cage monitoring coupled with AI quantification of phenotypes from video footage could obviate many traditional phenotyping paradigms that rely on animal handling and restraint, while also capturing animal behavior in a more ethologically relevant setting.

The Nachman panel is currently part of the Special Mouse Strain Resource at JAX, which provides the framework and setting for its long-term maintenance and external distribution. Several additional strains are maintained in Dr. Nachman’s private colony and we endeavor to import these additional strains to grow current holdings at JAX. Our team is currently engaged in efforts to expand the collection of tools and resources presented here, including de novo genome sequences, gene expression datasets, and the initiation of an outbred population founded from a subset of 4 Nachman strains. Together, the inbred Nachman strain panel, its planned derived populations, and accompanying ‘omics resources are poised to enable new discoveries in the basic, biomedical, and preclinical research spheres.

## MATERIALS AND METHODS

### Inbred Strain Development

Wild house mice were caught in 2013 from five geographic locations (Figure 1) using Sherman live traps baited with oats. To avoid catching closely related individuals, each mouse was trapped at least 500m from every other mouse. Mice were transported to UC Berkeley and held in quarantine. Mice from each geographic region were randomly paired to create inbred lines which were propagated through brother-sister mating for at least 10 generations. Initially, 10 independent lines were established from each location. No attempts were made to rescue lines that exhibited infertility due to inbreeding depression, ensuring that surviving lines likely harbor low deleterious mutation loads. All animal procedures were conducted under protocols approved by the UC Berkeley Animal Care and Use Committee. The founders of these strains were prepared as museum specimens and have been deposited in the collections of the UC Berkeley Museum of Vertebrate Zoology.

### Mouse Breeding, Rederivation, and Husbandry

Mice from 20 incipient inbred strains were imported from the Nachman Laboratory mouse colony at the University California Berkeley to the Jackson Laboratory Importation Facility (**Supplementary Table 2**). Strain colonies were expansion bred to obtain cohorts of >20 breeding-aged females for oocyte collection. To expedite breeding, strains were mated with some allowance for breeding between generations, a departure from strict sib-sib inbreeding.

Females were superovulated according to standard procedures (66) and harvested oocytes were *in vitro* fertilized with sperm from conspecific males (67). Embryos were cultured to the 2-cell or 8-cell stage before implantation into pseudopregnant QSi5/IanmTaftJ x C57BL/6J (JAX strains 027001 x 000664) F1 recipient dams of high health status. Live-born pups were transferred to the Dumont Laboratory colony where they were maintained at intermediate health status by strict sib-sib mating. A representative male and female from each successfully imported strain was genotyped on JAX’s standard 54-SNP panel to assure expected strain identity and safeguard against contamination with known laboratory stocks.

Mouse husbandry, euthanasia, and all animal procedures were conducted in compliance with an animal care protocol approved by The Jackson Laboratory’s Animal Care and Use Committee (Animal Use Summary # 18070). Mice were housed under SPF conditions in sterile plastic caging and subject to biweekly cage changes. All animals were fed 6% sterilized rodent chow (Lab Diet ® Formulation 5K0G) and provided with acidified water *ad libitium.* Cages were supplied with aspen bedding material and enriched with nestlets, crinkle paper nesting material, red igloos, and cedar blocks for gnawing.

### Sequencing

DNA extraction, library preparation, quality control, and sequencing were performed by the Genome Technologies Scientific Service at The Jackson Laboratory. An initial survey of short-read Illumina whole genome sequences from three strains (SARA/NachJ, SARB/NachJ, MANB/NachJ) pointed to high levels of structural variation relative to the mm39 reference genome (**Supplementary Figure 11**). This discovery motivated our use of long-read PacBio HiFi technology for all subsequent sequencing.

High molecular weight DNA was isolated from spleen tissue of a single male from each of the 11 imported Nachman wild-derived inbred strains using the Wizard DNA Purification Kit (Promega) or the Monarch HMW DNA kit (NEB) according to manufacturer’s instructions. DNA concentration and quality were assessed using the Nanodrop 2000 spectrophotometer (Thermo Scientific), the Qubit 3.0 dsDNA BR Assay (Thermo Scientific), and the Genomic DNA ScreenTape Analysis Assay (Agilent Technologies). DNA quality was assessed to be high (260/280 > 1.79 and 260/280 < 1.86, 260/230 > 1.99) and suitable for input for PacBio HiFi library construction.

PacBio HiFi libraries were constructed for each sample the SMRTbell Express Template Prep Kit 2.0 (Pacific Biosciences) according to the manufacturer’s protocols. Briefly, the protocol entails shearing DNA using the g-TUBE (Covaris), ligating PacBio specific barcoded adapters, and size selection on the Blue Pippin (Sage Science). The quality and concentration of the library were assessed using the Femto Pulse Genomic DNA 165 kb Kit (Agilent Technologies) and Qubit dsDNA HS Assay (ThermoFisher), respectively, according to the manufacturers’ instructions. The resultant library for each strain was sequenced on a single SMRT cell on the Sequel II platform (Pacific Biosciences) using a 30 hour movie time.

### Cytogenetic Chromosome Analysis

Spermatocyte cell spreads were prepared from testis tissue of males aged >8 weeks and immunostained as previously described (68). Antibodies used were a polyclonal antibody against mouse SYCP3 (1:100 dilution; Novus Biologicals, cat. # NB300-231) and human anti-centromere protein (1:100 dilution; Antibodies Incorporated, cat # 15-235). Cells were then imaged on a Leica DM6B upright fluorescent microscope equipped with fluorescent filters, LED illumination, and a cooled monochrome Leica DFC7000 GT 2.8 megapixel digital camera. Approximately 20 cells were captured per strain. Fluorescent intensity and background signal were manually adjusted, and the number of chromosomes counted using ImageJ (v 1.53k).

### Read Mapping and Variant Calling

Read quality and library coverage were assessed using *fastp* (69). Individual SMRT cells yielded between 22.04 and 38.04 Gb of unique sequence data, with an average read length of 12.9kb (**Supplementary Table 4**). Reads were then mapped to the GRCm39 reference sequence using minimap2 invoking the “HIFI” preset. Per sample single nucleotide calling was performed using DeepVariant (v1.2.0) under the “PACBIO” model. Per sample gVCF files were then merged using glnexus (v1.2.7) under the DeepVariantWGS configuration to produce a joint call set. Sites with missing data, genotype quality < 30, and indels were subsequently filtered using vcftools (v 0.1.16). We further eliminated sites with heterozygous calls as these sites are potentially enriched for false positives given our modest sequencing coverage. We refer to this callset as the “Nachman only” callset.

Variants in the Nachman strain panel were compared to those previously discovered in laboratory inbred strains in order to assess the extent to which this new resource captures novel genetic diversity. We downloaded GRCm39-mapped bam files for 51 inbred strains previously sequenced by the Mouse Genomes Project from the European Nucleotide Archive (PRJEB47108). Per sample variant calling was performed using the “WGS” model in DeepVariant (v1.2.0). Joint variant calling of laboratory strain genomes and the 11 inbred Nachman strains was carried out using glnexus (1.2.7) in DeepVariantWGS mode. As above, indels and variants with genotype quality < 30 were removed using bcftools (v 0.1.19). Sites with >10% missing data and heterozygous calls were also excluded. We refer to this callset as the “Sanger-Nachman” callset. Variant effects were predicted using the Variant Effect Predictor (Ensembl release_109.3) using the GRCm39 *Mus musculus* assembly.

Identical methodology was adopted to generate a DeepVariant joint call set for the Nachman strains and X wild-caught *M. m. domesticus*, Y *M. m. musculus*, Z *M. m. castaneus*, and A *M. spretus*. SRA accession numbers for the wild mouse genome sequences used are provided in **Supplementary Table 8**. We refer to this callset as the “wild-Nachman” callset. Again, sites with >10% missing data, genotype quality < 30, and indels were excluded using bcftools (v 0.1.19).

Basic statistics were computed on each call set using bcftools *stats*. Callsets were partitioned and intersected using *bcftools view* and and *bcftools isec*, respectively.

### Genome Sequence Analysis

SNP variation in the Sanger-Nachman callset was summarized via principal component analysis. We first eliminated six strains of non-*Mus musculus domesticus* ancestry from the callset (CAST/EiJ, CZECHII/EiJ, JF1/MsJ, MOLF/EiJ, PWK/PhJ, and SPRET/EiJ) and removed variants that were no longer segregating using the *view* command in bcftools (v 0.1.19) and the flags *-q 0.01 -s ^CAST_EiJ,CZECHII_EiJ,JF1_MsJ,MOLF_EiJ,PWK_PhJ,SPRET_EiJ*. Variants were greedily thinned to include only those with *r^2^* > 0.2 using the following command in plink (v1.90b6.18):

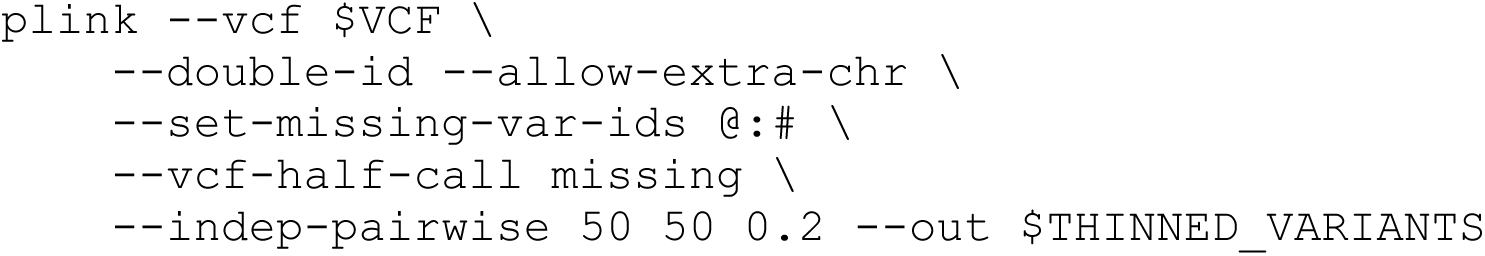

Principle component analysis was performed on the pruned set of variants using the –pca command in plink:

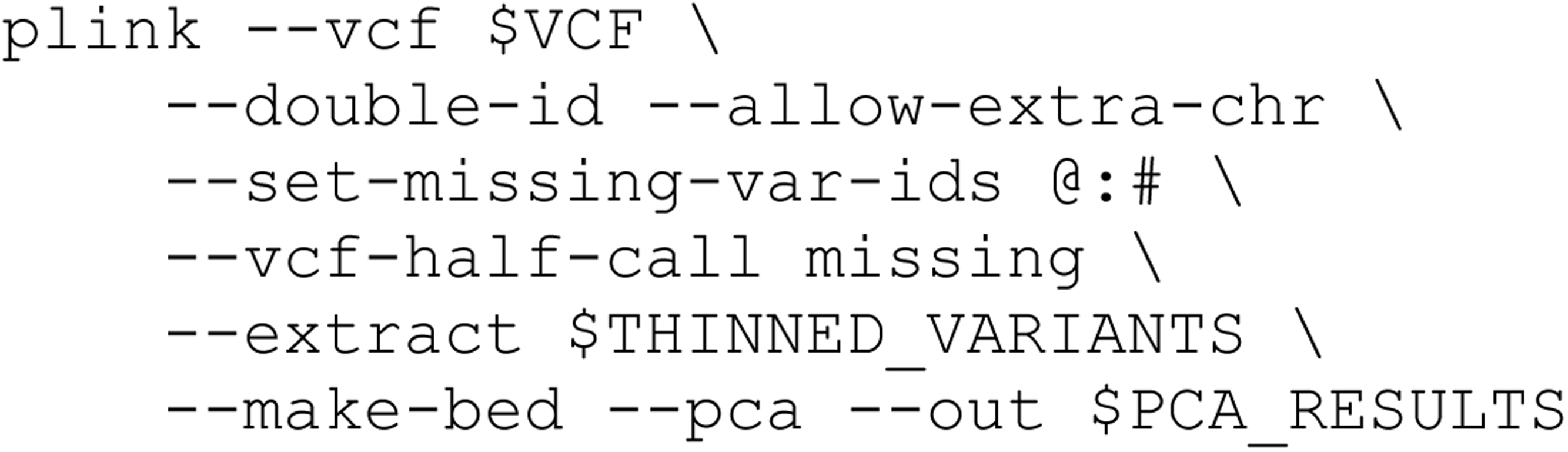

A maximum likelihood phylogenetic tree was constructed from the LD-thinned SNPs on chr19 in the Sanger-Nachman callset using *phyml* (version 3.3). We focus on chr19 for computational ease. We first created a fasta format file from the thinned chr19 SNPs in vcf format using a custom perl script (vcf_to_fasta.pl). The output file was then converted to phylip format using the SeqIO.write function in BioPython 1.43 library. A maximum likelihood tree was constructed under a GTR model of nucleotide evolution, with nucleotide frequencies computed from the empirical data, and with the transition/transversion ratio, proportion of invariant sites, and gamma distribution of rate classes estimated via maximum likelihood. The executed command was:

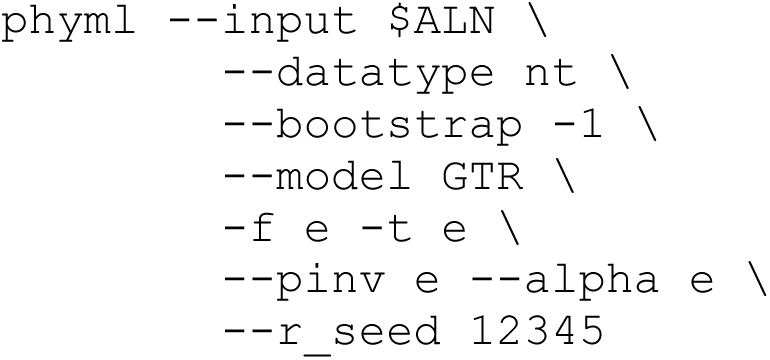

Identical methodology was used to perform PC analysis on the Nachman-wild callset. Prior to analysis, we first filtered the vcf file to include only wild *M. m. domesticus* samples, recognizing that PCA results would be dominated by subspecies-level diversity if samples from *M. m. musculus* and *M. m. castaneus* were retained. We also further subset the Nachman-wild callset to exclude *M. m. domesticus* samples from Iran.

### Introgression Analysis

Nachman strain genomes were scanned for potential signatures of introgression from *M. m. castaneus* and *M. m. musculus* using the allele sharing methods implemented in *Dsuite*. Wild *M. m. domesticus* and *M. m. castaneus* (or *M. m. musculus*) genomes in the wild-Nachman callset were utilized as P2 and P3, respectively, with *M. spretus* genomes specified as the outgroup taxon (**Supplementary Table 8**). *Dsuite Dtrio* was run separately on each chromosome and with each inbred Nachman strain individually profiled as P1. Chromosome-level statistics were integrated into genome-wide estimates using *Dcombine*. To pinpoint specific sites of introgression between *M. m. castanus* and each focal strain, a windowed analysis was performed using the *Dinvestigate* command in *Dsuite* with a window size of 5000 informative sites and 2500 site slide. Output statistics were plotted in RStudio (2022.02.0, Build 443). We focus on sites with an excess of derived alleles shared between each of our profiled Nachman lines and *M. m. castaneus*, and extracted the 5% of windows with the most extreme *f_d_* estimates consistent with potential introgression from *M. m. castaneus*. These outlier windows were then intersected with Ensembl genes (v. 109; *Mus musculus* GRCm39) using bedtools *isec* (v. 2.28.0) and subjected to a GO Enrichment Analysis using the online enrichment analysis tool on the GO Consortium Website (http://geneontology.org/). We specified the entire set of annotated genes in *Mus musculus* as background to identify specific biological processes enriched in putative regions of *M. m. castaneus* introgression. Enrichment was determined by a Fisher’s Exact test with False Discovery Rate of *P* < 0.05.

### Structural variant discovery and calling

We identified SVs in the 11 Nachman wild-derived inbred strain genomes using both *pbsv* and *sniffles2* (v. 2.0.7). *pbsv* was first run on each sample in *discover* mode to identify read signatures consistent with possible SVs. SVs where then called and samples jointly genotyped by executing pbsv in *call* mode. Tandem repeats in the GRCm39 assembly were identified using the findTandemRepeats.py script (https://github.com/PacificBiosciences/pbsv/commit/bcec7d382f3ea40158ed9cca3c5fef9686a76641) and supplied when executing sniffles to improve the accuracy of *sniffles2* calls in repetitive regions. Per sample SV calls generated by sniffles2 were merged and filtered to include only autosomal calls using *bcftools merge* (v1.10.2). Calls with close or overlapping breakpoints across samples were collapsed using *truvari* (v4.0.0), with the following parameters specified: -r 50 – pctsize 0.75 –pctovl 0.5 –pctseq 0.7 -s100 -k common. We then intersected *pbsv* and *sniffles2* SV calls using *truvari bench* to produce a higher confidence call set. We used the pbsv callset as the “truth” set and invoked the following command line parameters: -r 50 -pctsize 0.75 –pctovl 0.5 –pctseq 0 –dup-to-ins –passonly -s 100. SVs were annotated using the Ensembl Variant Effect Predictor (release 109.3) and annotations from the GRCm39 assembly.

Deletions and insertions in this intersection call set were converted to fasta format and input to RepeatMasker (v 4.1.5). The following repeatMasker command was executed: `RepeatMasker -e hmmer -q -species “mus musculus” -lcambig -nocut -div 50 -no_id.` To quantify the abundance of various transposable elements (TEs) within structural variants, we eliminate partial TEs, defined as those where <80% of the consensus transposable element sequence is represented within the structural variant.

Lastly, SV calls were intersected with a prior deletion callset released by the Sanger Mouse Genomes Project (https://ftp.ebi.ac.uk/pub/databases/mousegenomes/REL-1606-SV/REL-1606-SV/mgpv5.SV_deletions.bed.gz). Sanger call coordinates were first lifted over from mm10 to mm39 coordinates using the UCSC Genome Browser liftover tool. Shared deletions (>75% reciprocal overlap) were identified using *bedtools intersect* (v. 2.28.0).

### Animal Phenotyping

We developed a phenotyping pipeline loosely modeled on that used by the KOMP2 Project (https://www.mousephenotype.org/impress/index), with modifications to account for the overall health and wildness of the Nachman strains (**Figure 6**). Phenotypes broadly profile disease-relevant neurobehavioral, physiological, metabolic, biochemical, and morphological trait variation.

Mice were organized into 16 phenotyping cohorts ranging in size from 6-16 animals (mean = 13.4; **Supplementary Table 14**). With the exception of cohort 13, each cohort was composed of animals from multiple strains (N = 2-5 strains) and mice of both sexes. (Cohort 13 included only animals from strain GAIA/NachJ.) Ten of the 16 cohorts included C57BL/6J mice as controls (n = 2-5 mice per cohort), providing a means for ensuring stability of phenotype measurements across cohorts. A total of 215 mice (108 female, 107 male) were phenotyped from 9 of the 11 Nachman lines in two waves. The first wave included 5 cohorts that were phenotyped August-November 2020. The second wave included the remaining 11 cohorts which were phenotyped between June 2021 and October 2021. Strains EDMB/NachJ and TUCA/NachJ were not imported in time for inclusion in this phenotyping effort.

All live animal phenotyping was performed by trained staff in the Center for Biometric Analysis at The Jackson Laboratory (JAX). Animals were ∼4 weeks (± 6 days) at study intake and exited the phenotyping pipeline at 19 weeks (± 6 days). Terminal collections were performed by the Necropsy Scientific Service at JAX, with samples subsequently transferred to the Clinical Chemistry and Histopathological Sciences Scientific Services at JAX.

#### Body Composition Assessment by Nuclear Magnetic Resonance

Body composition analysis was carried out on mice at weeks 4, 8, 12, 16, and 19. Conscious, unrestrained mice were individually placed in an acrylic tube that was inserted into a Bruker Minispec LF50 Body Composition Analyzer to yield non-invasive estimates of mass and percentage estimates of fat, lean, and water mass.

#### Frailty Assessment (9 weeks)

Mice were gently restrained for visual assessment of 29 metrics of physical condition using a modified version of the protocol described in (56). Observational assessment of most parameters was qualitatively categorized as 0 if normal, 1 if highly abnormal, and 0.5 if intermediate. Exceptions include body temperature and body weight, which were measured in standard, qualitative units (Celsius and grams, respectively). A general frailty index score was computed by summing the scores for individual parameters (excluding body weight and body temperature).

#### Light Dark Test (10 weeks)

The light-dark test is a classic behavioral paradigm for quantifying anxiety-related behaviors in rodents (70). Following ∼60 minute acclimation to the testing room, mice were individually placed in a square, plexiglass arena (40 x 40 x 40 cm) divided into two compartments separated by a doorway. One side of the chamber was illuminated by an external light to ∼400-450 Lux, while the other side remained darkened. Mouse movements were tracked over a span of 10 minutes via an infrared photobeam 3-dimensional grid system invisible to the animals. Beam breaks were computationally decoded to provide information about latency to enter lighted side of the chamber, total number of transitions between the light and dark sides, total distance traveled, and time spent in the light versus dark sides.

#### Open Field Test (11 weeks)

Mice were first acclimated to the testing room for ∼60 minutes. Animals were individually placed into the center of a square plexiglass arena (40 x 40 x 40 cm) overlaid by a sensitive infrared photobeam three-dimensional grid. Beam breaks were computationally decoded into quantitative measurements of distance traveled, rearing, time spent within certain zones of the arena, and repetitive behaviors over the span of 60 minutes.

#### Spontaneous alternation with Y-maze (12 weeks)

Following a 60-minute room acclimation, a single mouse was placed into the center of an opaque Y-shaped testing arena composed of three equally sized arms each measuring 25-35 cm long, 5-6 cm wide, and with walls extending 12-18cm high. Mouse movements were recorded by a camera interfaced with tracking software (Noldus Ethnovision) to quantify distance traveled, arm entries, time spent in each arm, and the number of correct alternations (*i.e.,* a visit to each of the three arms before returning to any arms).

#### Home Cage Wheel Running (13 weeks)

Singly housed mice were provided with low profile running wheels (Med Associates) equipped with wireless transmitters for 3 days. Transmitters recorded the number of wheel revolutions and time on one-minute intervals. Data were forced onto a common timeline delimited by 18:00:00 on Day 1 to 6:00:00 on Day 3, converted to distances based on running wheel diameter (15.5cm), and aggregated over 5 minute, 30 minute, 1h, 6h, and 12 h intervals for analysis.

#### Indirect Calorimetry (16 weeks)

Mice were acclimated to single-housing for 5-7 days prior to transfer to Promethion Core cages for continuous high-definition respiratory monitoring over a 5-day span. Rates of oxygen consumption and carbon dioxide production were assessed using a built-in respirometry system, allowing calculation of overall energy expenditure using manufacturer software. Food and water intake were continuously monitored via high precision weight sensors mounted to the cage lid. Cages were overlaid with a grid of high sensitivity infrared beams to track animal activity and locomotion in continuous time.

#### Glucose Tolerance Testing

Animals were fasted for 4 hours, weighed, and restrained to make an angled incision at the tail for repeat blood collection. An initial blood drop was placed on a glucose test strip and a baseline blood glucose recorded using a handheld glucometer. Animals were then administered sterile glucose solution at a dose of 2g/kg of body weight via intra-peritoneal injection. Blood glucose levels were monitored at 15, 30, 60, 90, and 120 minutes post injection using single drops of blood obtained from the tail incision.

#### Unconscious Electrocardiogram and Gross Morphology

Mice were anesthetized by exposure to isoflurane gas in an enclosed chamber. Once unconscious, animal was removed from the chamber and placed ventral side up on an ECG platform, with continued gas delivery administered through a nose cone. Paralube ophthalmic ointment was placed on the eyes to prevent drying during testing and recovery. Single-use per-gelled self-adhesive silver/silver chloride ring recording electrodes (MVAP Medical Supplies, EMG electrodes) were adhered to the palmer surface of both front feet and the plantar surface of the left hind food. Leads were connected to each electrode and attached to a FE132 BIoAmp (AD Instruments) electrical signal amplifier which was fed into a PowerLab 4/35 digital-to-analog converter system and 4-channel recorder (AD Instruments) connected to a laptop computer with LabChart software. Core temperature was monitored and regulated via rectal probe physiology monitoring system and automatic heat pad. The ECG tracing was recorded until 30 seconds of waveform signal was obtained.

While animals remained unconscious, overall length, body length, and tail length were measured using handheld calipers.

#### Terminal Collections

Approximately ∼400ul whole blood was collected from the submental vein of fasted mice (4 hour). Animals were then euthanized via CO_2_ asphyxiation in accordance with recommendations from the American Veterinary Medical Association. Blood samples were transferred to the Clinical Chemistry Scientific Service at The Jackson Laboratory for measurement of the following blood-based clinical traits: albumin, alkaline phosphatase, alanine transaminase, aspartate transaminase, blood urea nitrogen, enzymatic creatinine, glucose, total cholesterol, HDLD cholesterol, triglycerides, insulin, leptin, iron, total bilirubin, total protein, and complete blood count with differential. The following organs were carefully dissected from each animal by skilled staff in the Jackson Laboratory Necropsy Scientific Service: skeletal muscle, gonadal fat pad, brown adipose from between the shoulder blades, tail, skin (with hair), femur, spleen, kidneys, liver, brain, gonads, heart, lungs, eye. Tissues were weighed, frozen, and fixed in 10% NBF and paraffin embedded according to **Supplementary Table 30**.

#### Data Integration, Statistical Analysis, and broad sense heritability estimation

All phenotype data were wrangled into a single file in RStudio (v. 2022.02.0 Build 443) using the dplyr, tidyverse, data.table, readxl, and lubridate packages (**Supplementary Table 21**). We tested for overall strain effects on each phenotype using non-parametric Kruskall Wallis (**Supplementary Table 16**) and one-way ANOVA (phenotype value ∼ strain; **Supplementary Table 17**). Categorical traits collected as part of the frailty analysis (n = 27) showed little variation among strains (**Supplementary Table 22**) and were excluded from statistical analyses. All remaining phenotypes were continuous. Although the ANOVA assumption of normality does not hold for many analyzed traits, there is strong agreement of *P*-values estimated by these two statistical methods (Spearman’s Rho = 0.94, P < 2.2 x 10^-16^). Given that our objective is to flag phenotypes that vary across the Nachman panel, reported *P*-values are not corrected for multiple testing.

Broad sense heritability was estimated from one-way ANOVA output by the intraclass correlation method (71):

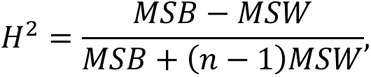

where MSB and MSW are the mean squares between and within strains, respectively. Heritability estimates were computed from one-way ANOVA tests run on each sex separately, as well as both sexes combined. Two-way ANOVA was also performed to test the impact of strain, sex, and their interaction on the variability observed in each trait (phenotype value ∼ strain * sex; **Supplementary Table 20**). Spearman’s rank correlate was used to assess trait correlations at the strain, strain x sex, and individual levels.

In parallel to this approach, we also fit a series of nested linear mixed effect models to assess the impact of strain, sex, and strain x sex interaction on each phenotype. Specifically, for each trait *t*, we fit the following series of models using the *lmer* command within the *lme4* (version 1.1-32) R package:

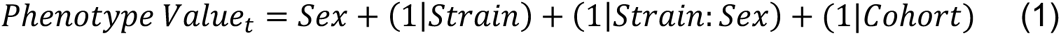

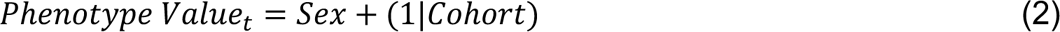

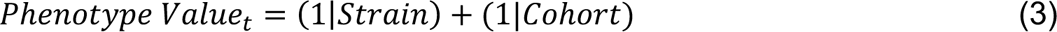

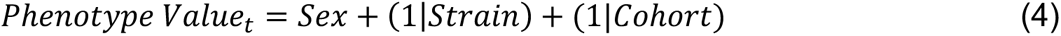

Sex was treated as a fixed effect, with strain, cohort, and the interaction between strain and sex treated as random effects. Nested models were compared by a log likelihood test using the *anova* function call in base R. Specifically, models 1 and 2 were compared to assess whether inclusion of strain as a random effect significantly improved model fit. Models 1 and 3 were compared to assess whether inclusion of sex significantly improved model fit, and models 1 and 4 were compared to determine whether the interaction between strain and sex meaningfully improved model fit. Owing to the modest amount of data and the complexity of these models, many models were singular, necessitating caution in the interpretation of results from this model fitting approach. Of the 1092 traits analyzed under this mixed model framework, only 405 provided non-singular model fits (**Supplementary Table 18**).

Many phenotypes are highly correlated with body weight (**Supplementary Table 15**), motivating us to also consider body weight adjusted trait values in our models. We fit the residuals from a simple linear regression of each trait value on body weight at 18 weeks in each of the models above. This had a similar impact on the number of singular model fits (1083 analyzed traits, with 446 with non-singular fits across models 1-4; **Supplementary Table 19**). We excluded body weight traits at all assessed timepoints, accounting for the reduced number of phenotypes analyzed under this framework.

#### Comparisons with MPD data

We accessed the CGDpheno1, CGDpheno2, and JAXKOMP-EAP datasets from the Mouse Phenome Database (72). These strain survey datasets were prioritized because: (1) they share multiple phenotypes in common with those measured in the Nachman strains, (2) they were generated within the CBA at JAX, providing methodological consistency in data collection and similarity in housing conditions, and (3) animals were not subject to any experimental treatments. Public datasets were combined with the Nachman data using base R commands and dplyr (v. 1.1.1), with attention paid to consistency of measurement units. Where needed, values were converted onto common unit scales using the appropriate conversion factors or mathematical manipulations.

## ACKNOWLEDGEMENTS

We thank Gabriela Heyer, Ketki Samel, Henry Thomas, Noelle Bittner, Felipe M. Martins, Sarah Banker, Brett Haines, and Kristin Lapointe for assistance with animal husbandry and breeding. We gratefully acknowledge the contribution of the Genome Technologies, Histopathology, Necropsy, Clinical Chemistry Scientific Services at The Jackson Laboratory for their contributions to this work. We also thank members of the Center for Biometric Analysis for their critical contributions toward phenotype data collection.

Importation, whole-genome sequencing, and phenotyping were supported by a JAX Director’s Innovation Fund Award to B.L.D., G.A.C., N.R., C.L, and J.W.

B.L.D. is supported by a Maximizing Investigators’ Research Award from the National Institute of General Medical Sciences (R35 GM133415). MWN was supported by RO1 GM074245, R01 GM127468, and R35 GM149304 from the National Institute of General Medical Sciences.

---

**Supplementary Figure 1.**
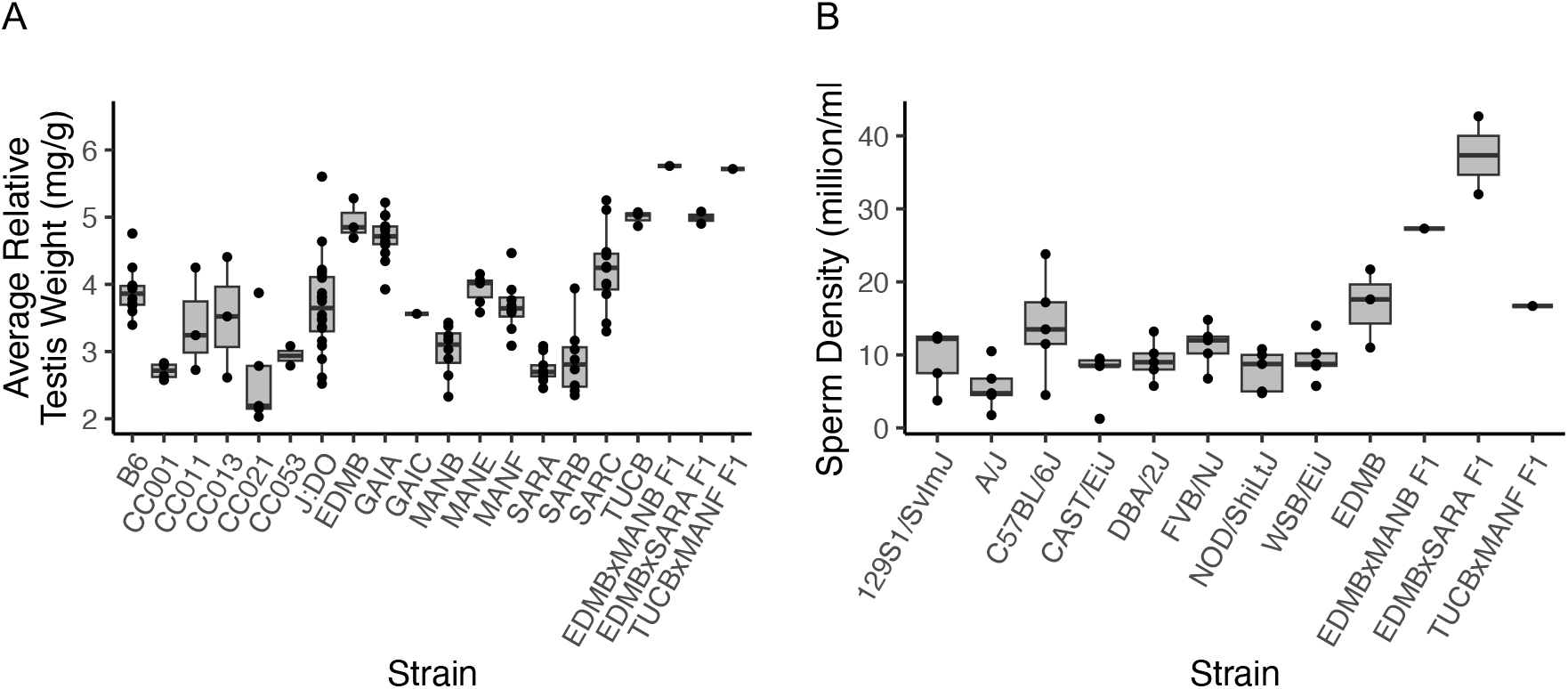
Male reproductive traits in the inbred Nachman strains, a subset of their F1 hybrids, and representative inbred strains. (A) Testis weight standardized by body weight for inbred Nachman strains and three derivative F1 hybrids. Values for the classical inbred strain C57BL/6J (B6), a sample of DO animals, and a representative subset of CC strains are included for comparison. (B) Sperm density estimates for 8 genetically diverse inbred mouse strains, EDMB, and 3 F1 hybrids derived from crosses between Nachman strains. Inbred strain sperm density estimates are from the Handel1 dataset on the Mouse Phenome Database.

**Supplementary Figure 2.**
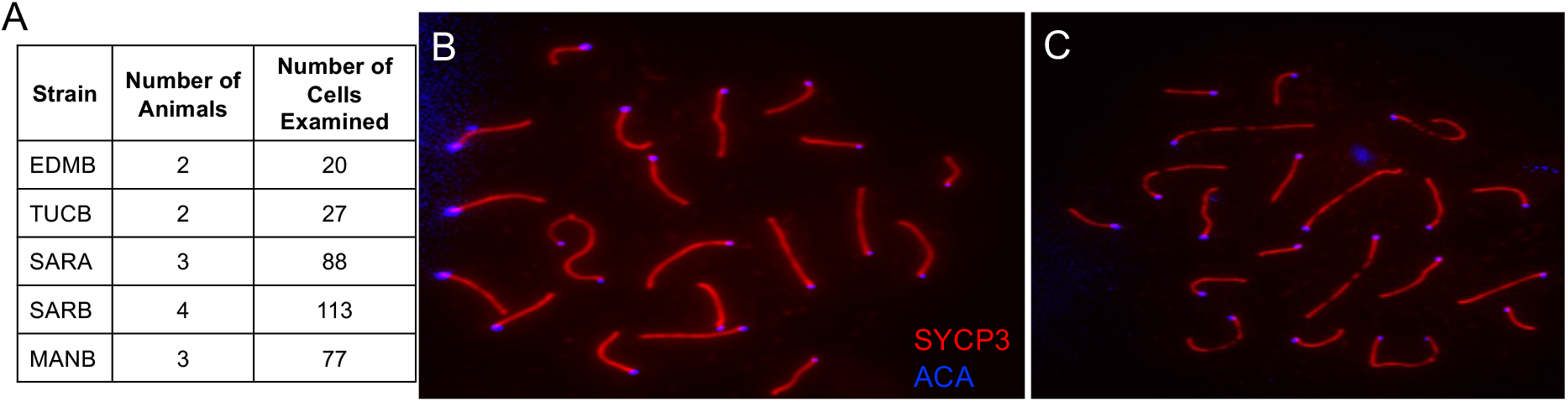
Cytogenetic analysis of meiotic spermatocyte spreads confirms 2N=40 karyotypes in Nachman strains. (A) Table summary of the number of individuals and number of cells analyzed. All cells derive from males and all harbor the standard house mouse karyotype defined by 19 acrocentric autosomal pairs and a pair of sex chromosomes. Representative pachytene-stage spermatocyte cell spreads from TUCB/NachJ (B) and EDMB/NachJ (C) stained with antibodies against SYCP3 (red), a component of the meiotic synaptonemal complex that aligns along the paired chromosome axes, and anti-centromere antibodies (blue).

**Supplementary Figure 3.**
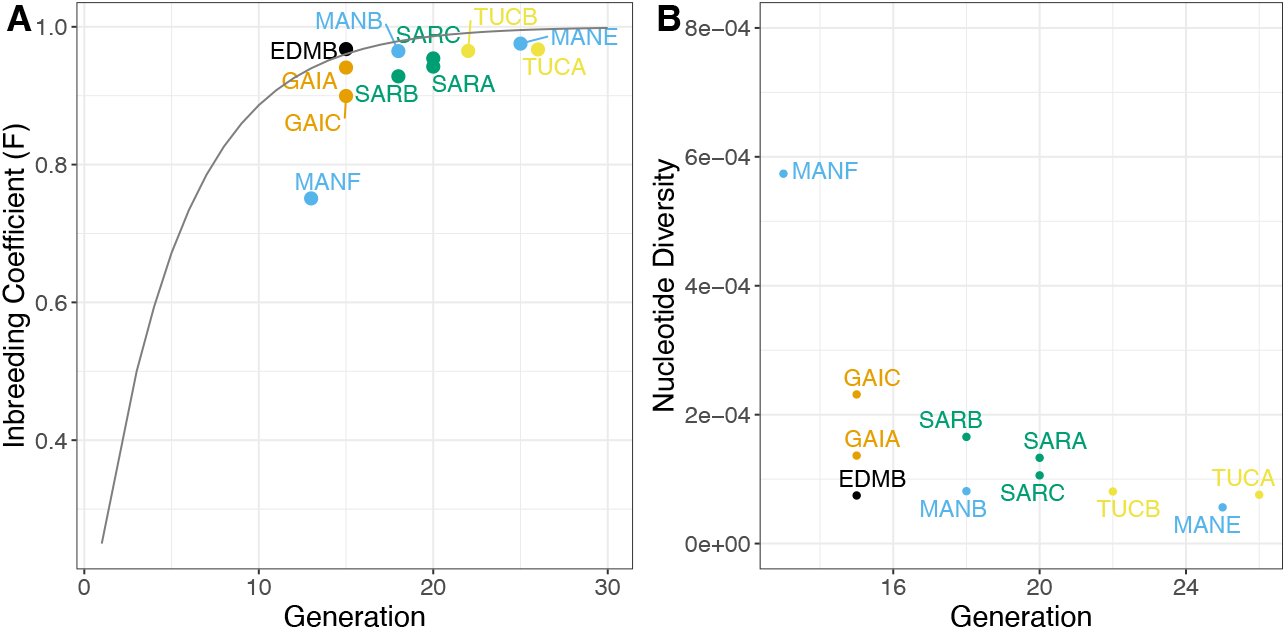
Heterozygosity declines as a function of inbreeding generation. (A) The observed inbreeding coefficient, *F*, was calculated using the method of moments estimator implemented in vcftools (v 0.1.16) (11,44,45). For most strains, the empirical *F* estimate closely tracks with the theoretical expectation for increasing inbreeding generation number (black line). Strain MANF/NachJ presents an exception, and is likely attributable to departure from strict sib-sib mating during the colony expansion effort that preceded strain rederivation. Per sample nucleotide diversity (*π*; B) declines with increasing inbreeding generation, as expected.

**Supplementary Figure 4.**
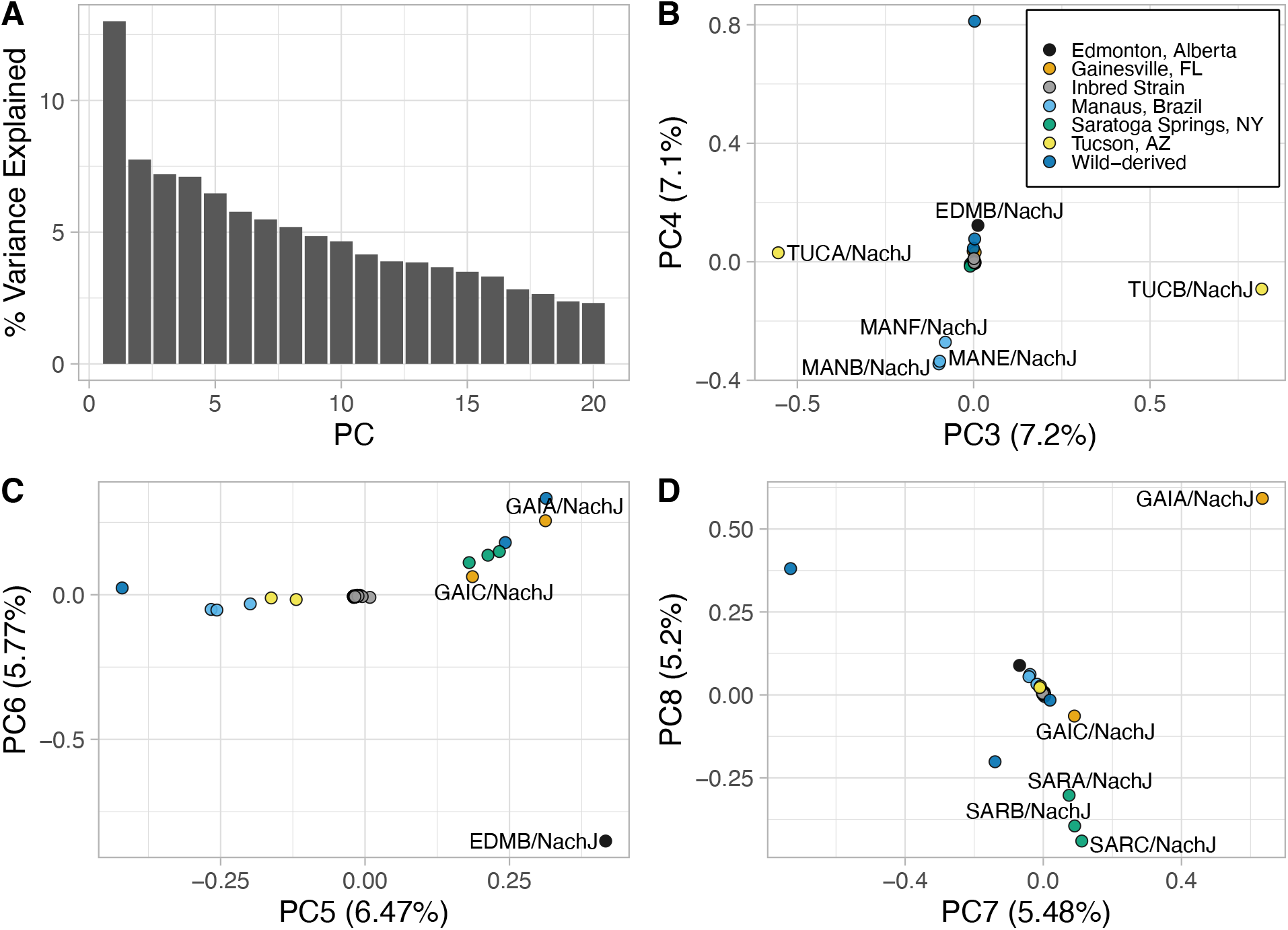
Higher principal component dimensions provide additional refinement to the genetic relationships between the Nachman and classical inbred mouse strains. (A) Percentage of total variance explained by each PC dimension 1-20. (B) PC3 versus PC4. (C) PC5 versus PC6. (D) PC7 versus PC8.

**Supplementary Figure 5.**
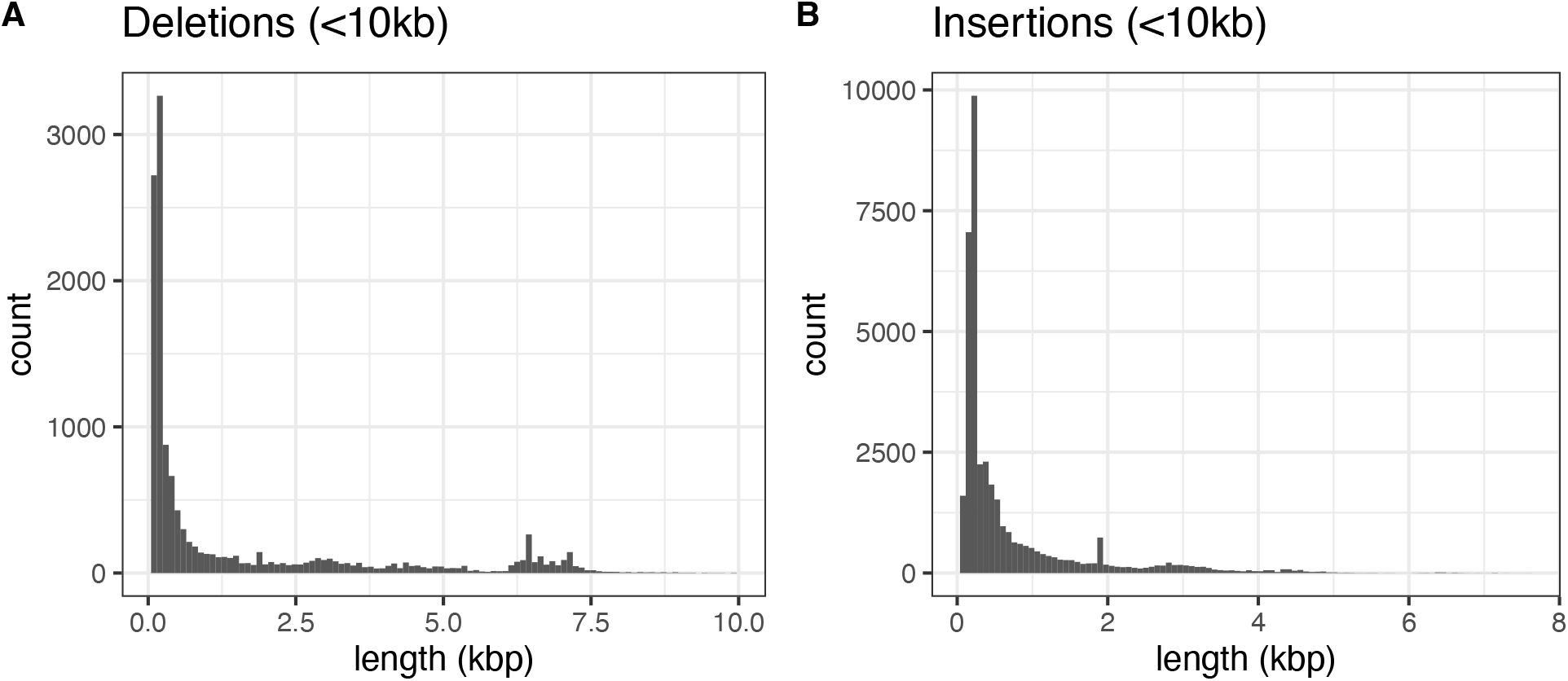
Distribution of deletion (A) and insertion (B) lengths across the Nachman strains. For ease of visualization, only SVs <10kb in length are plotted. Only 49 deletions and 1 insertion call exceed this cutoff.

**Supplementary Figure 6.**
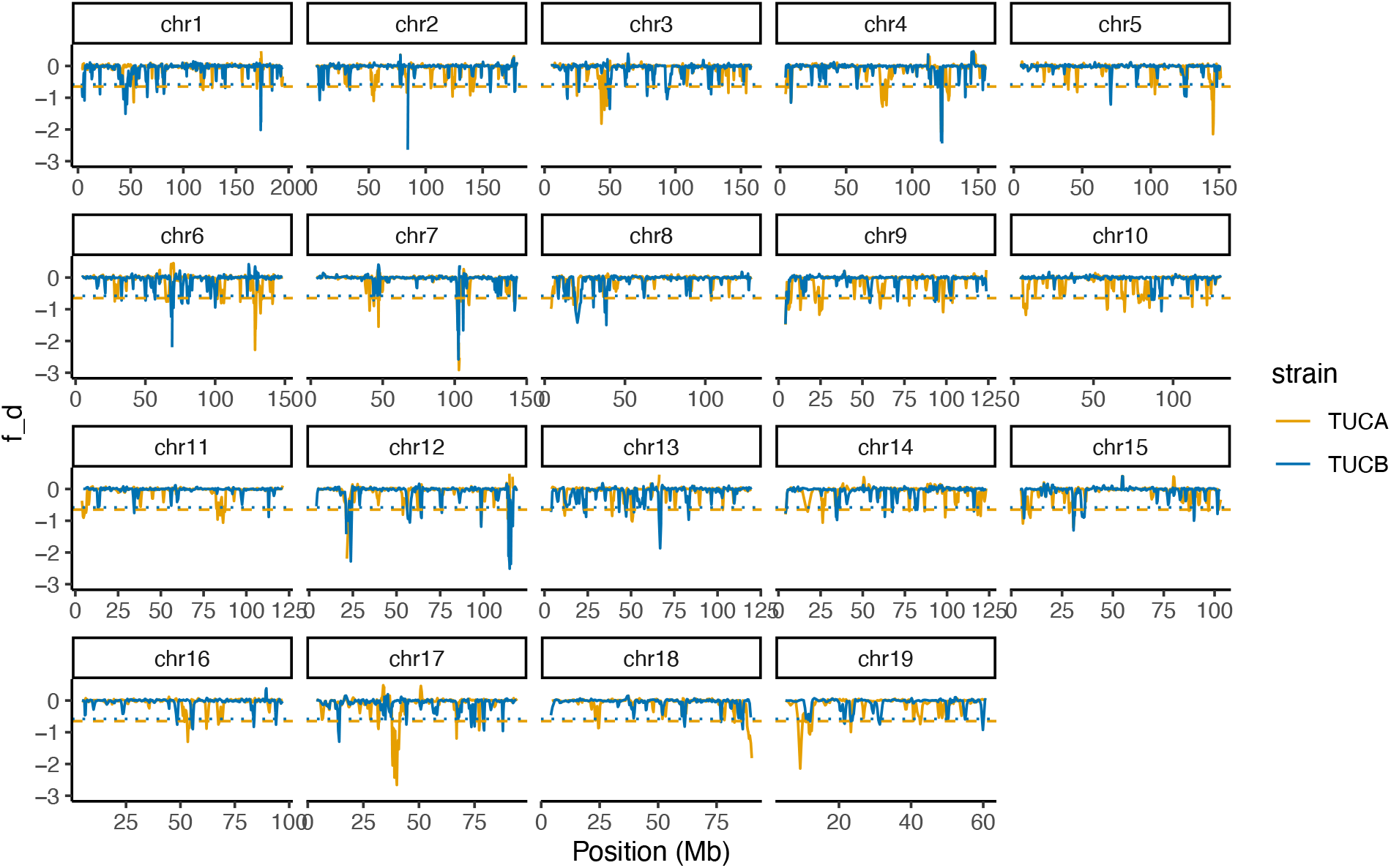
Genomic distribution of *f_d_* in TUCA and TUCB. *f_d_* was calculated in 5000 SNP windows, with a 2500 SNPs slide using genome sequences from wild-caught *M. m. domesticus*, wild-caught *M. m. castaneus*, and *M. spretus* (Supplementary Table 9). Dashed lines correspond to the 95th percentile of the most extremely negative *f_d_* statistics for each strain. The y-axis of each plot was clipped to show only values >-3 and <1 for ease of visualization.

**Supplementary Figure 7.**
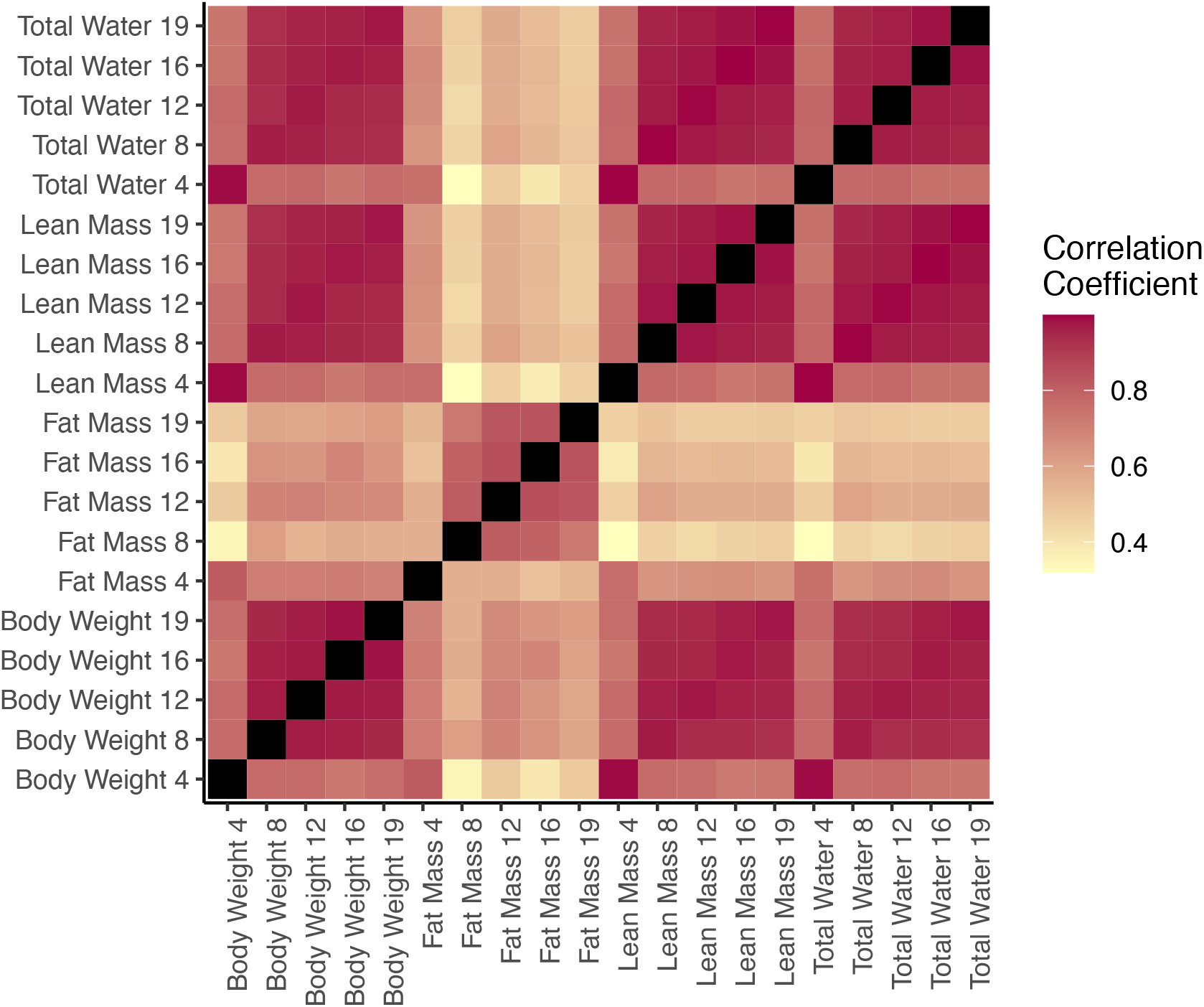
Correlation heat map for body composition and body weight metrics assayed at 4, 8, 12, 16, and 19 weeks of age. The magnitude of the correlation coefficient (Spearman’s rho) is indicated by the scale bar, with higher correlations denoted by darker red colors, and weaker correlations indicated in yellow.

**Supplementary Figure 8.**
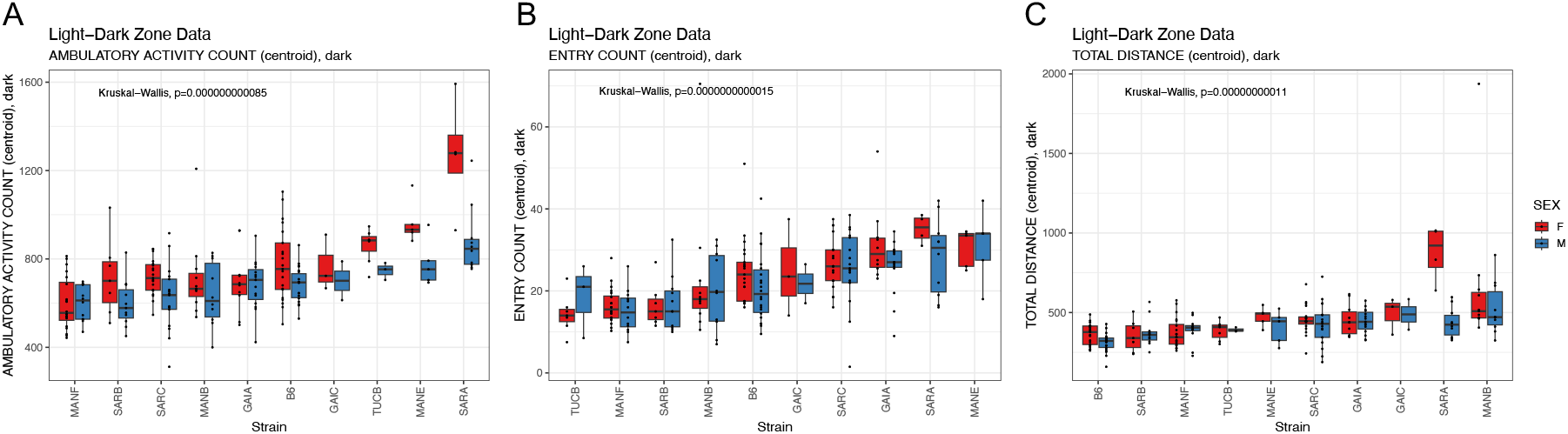
Strain variation in several anxiety-related metrics assessed by the light-dark test. In (C), distance is plotted on the y-axis in meters. The color legend presented in (C) applies to all panels.

**Supplementary Figure 9.**
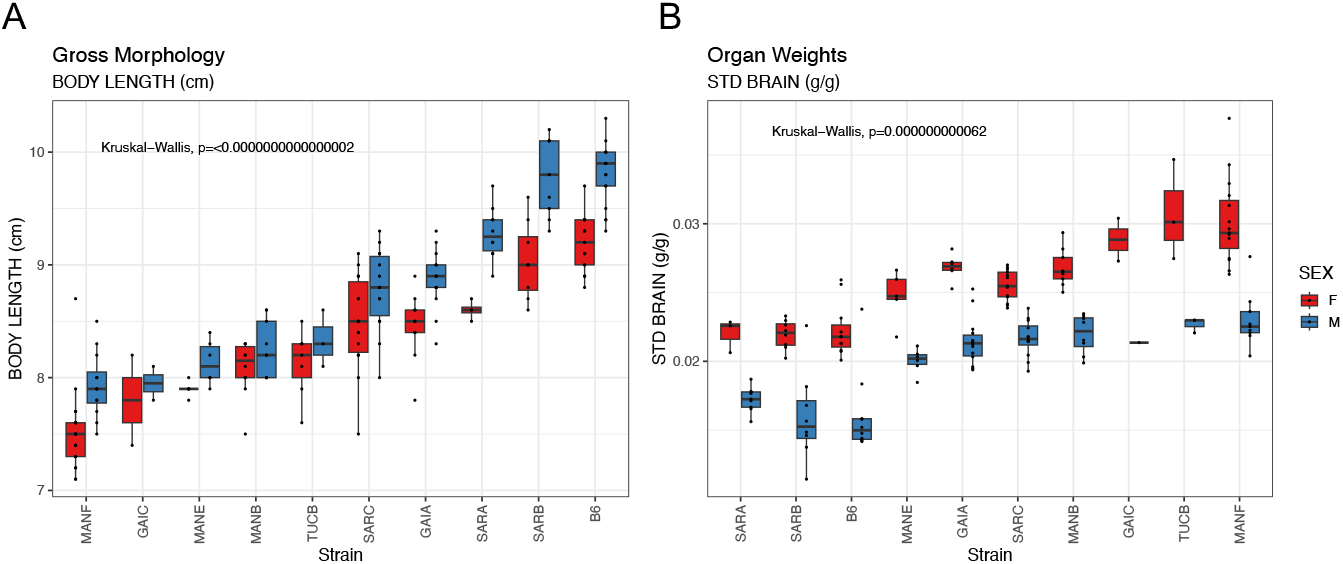
Strain and sex variation in body length (A) and (B) brain weight, standardized by total body weight.

**Supplementary Figure 10.**
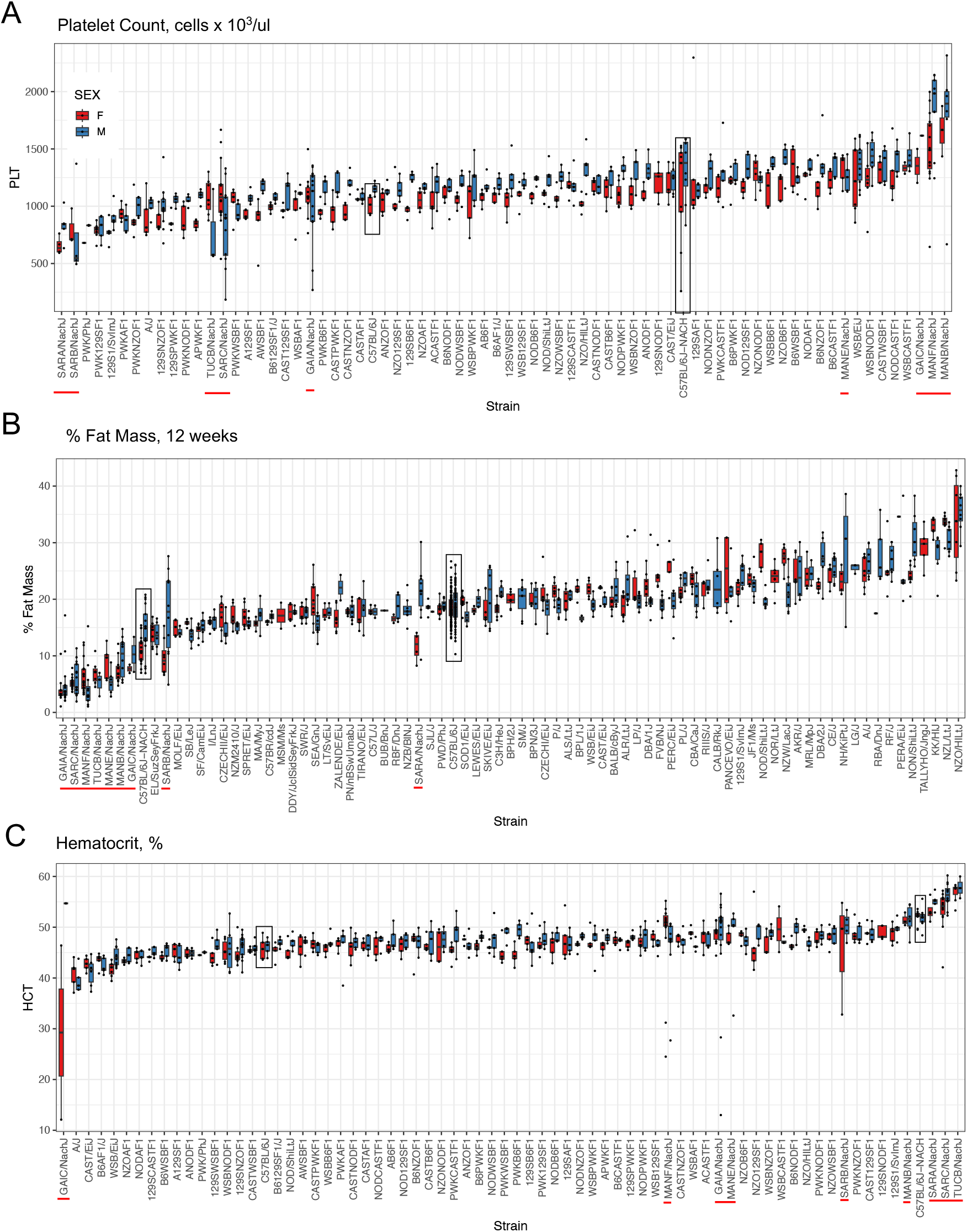
Nachman strains expand the range of phenotypic trait variance observed in current inbred strain and mouse diversity panels. (A) Platelet counts (cells x 10^3^/ul) for the Nachman strains were integrated with the CGDpheno3 dataset from the Mouse Phenome Database. (B) Percent fat mass at 12 weeks in the Nachman lines and strains profiled in the CGDpheno1 dataset. (C) Percent hematocrit in the Nachman lines and strains in CGDpheno3. The color legend in (A) applies to all panels. In each panel, Nachman strains are indicated by red bars along the x-axis and boxplots for C57BL/6J mice are indicated by black boxes. C57BL/6J mice phenotyped as controls alongside the Nachman strains are denoted as “C57BL/6J-NACH”; C57BL/6J animals phenotyped in other studies are indicated as “C57BL/6J” along the x-axis.

**Supplementary Figure 11.**
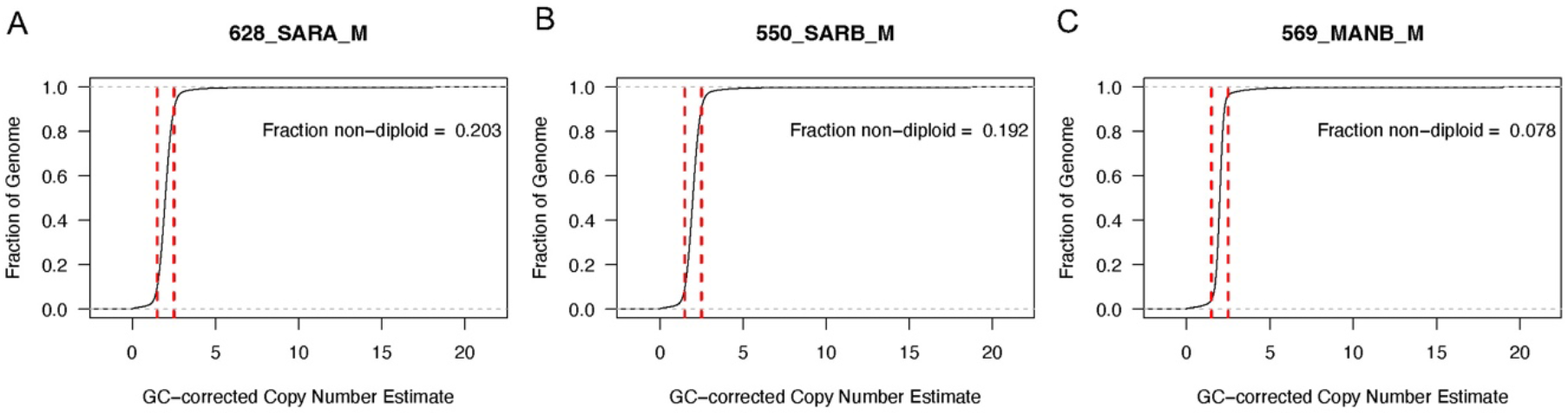
Cumulative distribution of copy number state in 1kb sliding windows across the autosomal genome for (A) SARA/NachJ, (B) SARB/NachJ, and (C) MANB/NachJ. To estimate copy number, read depth was first computed in 1kb sliding windows (no overlap) using *mosdepth* (executed command: mosdepth -n -b 1000 -t 2 -x $PREFIX $BAM). Raw read depth values were then corrected for potential GC-biases introduced during library preparation. Briefly, we used the mm39 reference genome to compute the observed GC content of each 1000bp window. GC content values were rounded to the nearest 0.001 and regions with identical GC content were binned. For each strain, we then computed the mean read depth across all genomic windows that fell into each GC content bin. Next, we fitted a second degree polynomial to the relationship between read depth and GC content using the scatter.smooth function in R and with span parameter of 0.7. For each GC-bin, we then computed the difference between the fitted polynomial and the genome-wide average read depth. These values correspond to the magnitude of “inflation” or “deflation” in read depth across windows of a given GC-content due to systematic GC biases in the data. The read depth value in each 1000bp window was then adjusted by the appropriate GC correction factor. Finally, these GC-corrected read depths were divided by the average per-sample coverage to convert into absolute copy number estimates. The cumulative distribution of autosomal copy number estimates was then computed using the *ecdf* function call within the *stats* package in Rstudio (version 2022.02.0, Build 443). The proportion of windows with copy number <1.5 and >2.5 (red vertical lines) was calculated as a proxy for the extent of structural variation in the strain genome relative to mm39.

